# CryoDataBot: a pipeline to curate cryoEM datasets for AI-driven structural biology

**DOI:** 10.1101/2025.09.09.675185

**Authors:** Qibo Xu, Leon Wu, Michael Rebelo, Shi Feng, Xinye Yu, Farhanaz Farheen, Daisuke Kihara, Z. Hong Zhou

**Affiliations:** California NanoSystems Institute, University of California, Los Angeles, CA 90095, USA; Department of Microbiology, Immunology, and Molecular Genetics, University of California, Los Angeles, CA 90095, USA; Department of Bioengineering, University of California, Los Angeles, CA 90095, USA; Department of Chemistry and Biochemistry, University of California, Los Angeles, CA 90095, USA; Department of Computer Science, Purdue University, West Lafayette, IN 47907, USA; Department of Biological Sciences, Purdue University, West Lafayette, IN 47907, USA

## Abstract

Cryogenic electron microscopy (cryoEM) has revolutionized structural biology by enabling atomic-resolution visualization of biomacromolecules. To automate atomic model building from cryoEM maps, artificial intelligence (AI) methods have emerged as powerful tools. Although high-quality, task-specific datasets play a critical role in AI-based modeling, assembling such resources often requires considerable effort and domain expertise. We present CryoDataBot, an automated pipeline that addresses this gap. It streamlines data retrieval, preprocessing, and labeling, with fine-grained quality control and flexible customization, enabling efficient generation of robust datasets. CryoDataBot’s effectiveness is demonstrated through improved training efficiency in U-Net models and rapid, effective retraining of CryoREAD, a widely used RNA modeling tool. By simplifying the workflow and offering customizable quality control, CryoDataBot enables researchers to easily tailor dataset construction to the specific objectives of their models, while ensuring high data quality and reducing manual workload. This flexibility supports a wide range of applications in AI-driven structural biology.

## Introduction

Accurately determining the three-dimensional structure of biomacromolecules is fundamental to understanding their function and guiding therapeutic development^1^. Traditional approaches like X-ray crystallography and nuclear magnetic resonance (NMR) spectroscopy have long been central to structural biology. However, these methods are often limited by poor protein solubility, sample heterogeneity, and intrinsic resolution constraints. Furthermore, they typically require considerable manual effort, such as sample crystallization. In contrast, cryogenic electron microscopy (cryoEM) enables high-resolution imaging of large and heterogeneous macromolecular complexes in their native conformations^2–6^. This capability allows cryoEM to reveal intricate structural details across diverse conformational states under physiological conditions.

Despite substantial advancements in cryoEM instrumentation and image processing, structure modeling remains a major bottleneck. Like crystallography and NMR, deriving atomic models from cryoEM data can take several weeks or even months and remains error-prone^7,8^, even with sophisticated tools such as PHENIX^9^, Coot^10^ and ChimeraX^11^. As cryoEM continues to expand in scope and target increasingly complex biomacromolecules, recent research has turned to artificial intelligence (AI) for optimizing cryoEM maps and automating modeling^12–18^. These methods often employ architectures such as convolutional neural networks, graph neural networks, and U-Net models^19–21^. Regardless of implementation, these approaches critically depend on the availability of high-quality training datasets characterized by structural diversity, accurate labels, and low redundancy^22^.

AI-based modeling tools such as DeepTracer^12^, ModelAngelo^13^, and CryoREAD^14^ typically build training datasets by retrieving cryoEM maps and associated atomic models from the Electron Microscopy Data Bank (EMDB)^23^ and the Protein Data Bank (PDB)^24^. These maps are then resampled to a uniform voxel size, and structural labels are generated from the atomic coordinates. The data, comprising resampled maps and labels, are partitioned into smaller 3D sub-volumes to support efficient model training. Some pipelines include additional quality enhancement steps, such as map–model fitness (MMF) evaluation and redundancy handling (Table 1). While these practices improve dataset fidelity and modeling accuracy, the absence of standardized construction pipelines across tools impedes reproducibility and complicates cross-comparisons. Moreover, the processed datasets are generally not publicly available, presenting a major hurdle for new developers who must rebuild training data from scratch.

**Table 1:** Comparison of training datasets used by existing AI-driven cryoEM modeling tools, Cryo2StructData dataset, and CryoDataBot-generated dataset. Notably, while some tools have implemented quality enhancement procedures, their datasets exhibit non-uniform voxel sizes and remain unavailable to the public. Cryo2StructData offers a large, accessible dataset, but omits several quality control steps used by other software. CryoDataBot provides users with flexibility by enabling fully customizable and quality-controlled dataset generation.

| Software |  | DeepTracer | ModelAngelo | CryoREAD | Cryo2StructData | CryoDataBot |
| --- | --- | --- | --- | --- | --- | --- |
| Dataset Overview | Macromolecule Type | protein | protein, RNA & DNA | RNA & DNA | protein | Customized* |
|  | Entries (Initial) | 1,800 | 3,715 | 1,384 | 7,600 | Customized |
|  | Entries (Curated) | 1,800 | ~ 700 | 290 | 7,600 | Customized |
|  | Resample Voxel Size | 0.5 Å | 1.5 Å | 1.0 Å | 1.0 Å | Customized |
| Quality Enhancement Procedures | MMF Evaluation | No | Yes | Yes | No | Yes |
|  | Redundancy Handling | No | Yes** | Yes | No | Yes |
| Public Availability |  | No | No | No | Yes | Yes |

\* user-defined; can include proteins, DNA, RNA, or specific macromolecular types such as ribosomes, spliceosomes, etc.
\*\* manual work involved

A recent effort, Cryo2StructData^25^, began to address these issues by releasing a large-scale, publicly available cryoEM dataset with curated labels and consistent formatting, providing a valuable resource for the development and benchmarking of AI-based modeling tools. Despite its strengths, Cryo2StructData leaves room for further refinement in several key areas that impact downstream usability. Notably, the absence of quality control procedures—such as map–model fitness (MMF) evaluation and redundancy filtering (Table 1)—may necessitate additional data refinement prior to training AI models, in order to ensure consistency and reliability across downstream tasks. Additionally, the use of a fixed voxel resolution (1.0 Å) also limits compatibility with diverse AI architectures, which often require tailored input scales (e.g., 0.5 Å for DeepTracer^26^, 1.5 Å for ModelAngelo^13^). Addressing these limitations through flexible resolution support and integrated quality metrics would broaden the dataset’s applicability.

To overcome current limitations in cryoEM dataset preparation for AI applications, we developed CryoDataBot—a GUI-based pipeline that enables researchers to generate customized, high-quality datasets with minimal input. It supports filtering based on residue-level similarity, Q-scores^27^, MMF thresholds, and EMDB metadata^28^. By offering a standardized and modular framework, CryoDataBot streamlines dataset construction, facilitates targeted model training, enhances workflow reproducibility, and empowers researchers in AI-driven structural biology.

## Results

### Pipeline of CryoDataBot and Functionality of Each Module

We design CryoDataBot as a tool to automate generation of high-quality datasets for AI-driven atomic structure modeling. First, it should provide a simple interface for users to enter query parameters and seamlessly communicate with the Electron Microscopy Data Bank (EMDB), extract and parse relevant metadata, apply rigorous quality control measures, and retrieve associated structural data—cryoEM maps and atomic models. Second, the tool should generate accurate structural labels and output fully formatted datasets suitable for training and evaluating deep learning models in macromolecular structure prediction. Figure 1 provides an overview of the implemented data processing pipeline of CryoDataBot, which consists of four major modules: Metadata Collection, Metadata Curation, Structural Data Conditioning, and Customized Dataset Construction, along with an illustration of the graphical user interface (GUI).

**Fig. 1:**
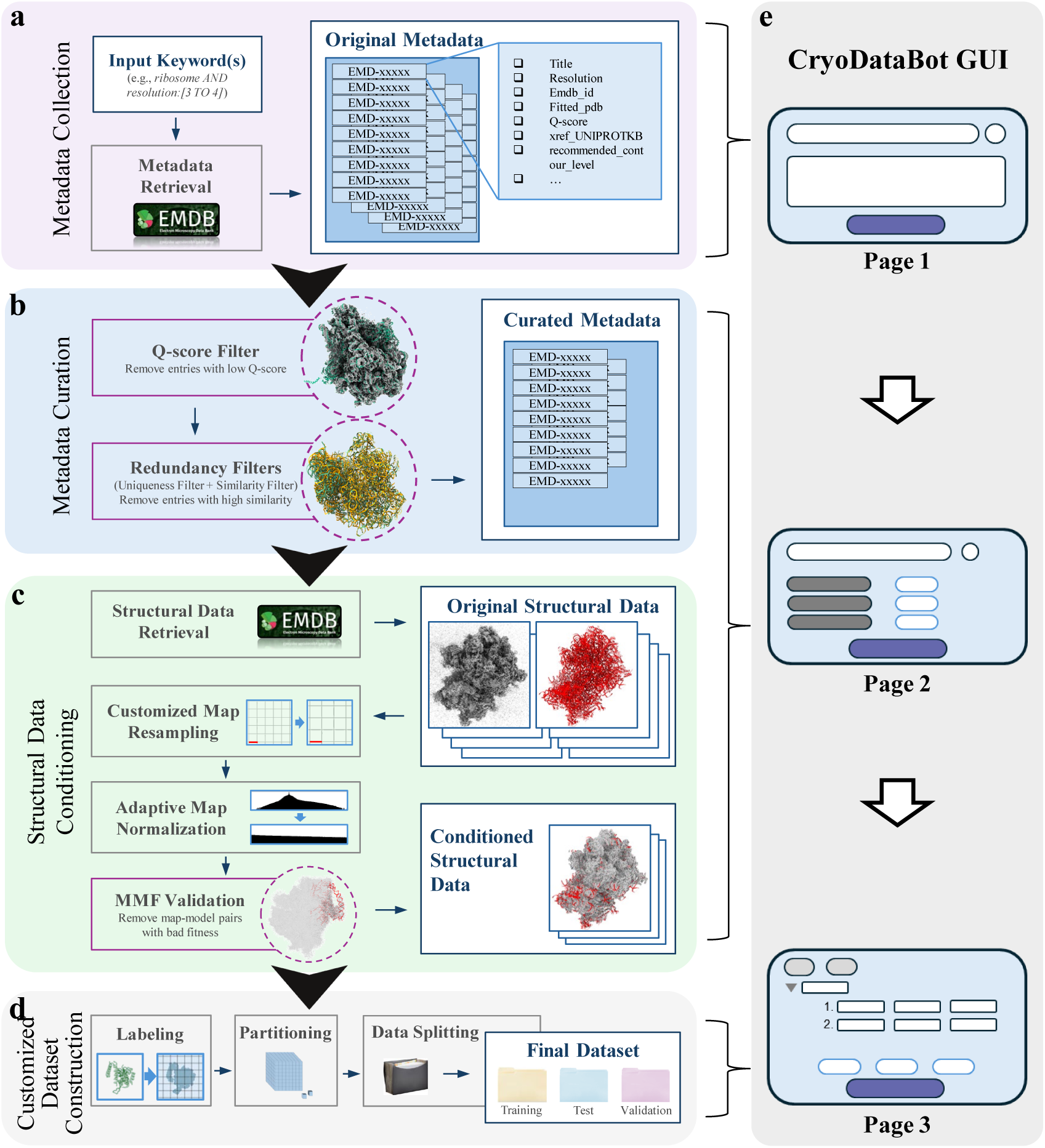
Pipeline of CryoDataBot. CryoDataBot consists of four functional modules. **a**, Metadata Collection: entries are retrieved from EMDB using user-defined keywords. **b**, Metadata Curation: metadata undergo a multi-stage quality control pipeline to ensure map–model fitness and reduce overall redundancy. **c**, Structural Data Conditioning: cryoEM maps and atomic models are retrieved, resampled to a customized uniform voxel size, and normalized to a 0–1 density range using recommended contour levels for denoising. Map–model fitness is then validated to ensure the accuracy and completeness of the atomic models. **d**, Customized Dataset Construction: conditioned structural data are labeled based on user-specified atom types, partitioned into 3D sub-volumes, and split into training, validation, and test sets tailored for AI applications. For further details, see Methods. **e**, Cartoon representation of the graphical user interface (GUI) of CryoDataBot. The first page of the GUI corresponds to module **a**, the second page covers modules **b** and **c**, and the third page is dedicated to module **d**.

The **Metadata Collection** module facilitates the gathering of raw data (Fig. 1a). It starts by querying the EMDB using a list of user-defined keywords and then automatically collects relevant metadata (Methods). The collected metadata include fields such as entry titles, EMDB IDs, fitted PDB IDs, Q-scores, cross-references to the Universal Protein Knowledgebase (UniProtKB) and AlphaFold, recommended contour levels, and other relevant annotations. To ensure completeness and reduce redundancy, entries missing critical fields (e.g., fitted PDB ID) or containing duplicate EMDB IDs or titles are excluded from the original metadata.

Through the **Metadata Curation** module, the original metadata are subjected to a multi-stage quality control pipeline (Fig. 1b). The first criterion is the Q-score, which quantitatively assesses the atom-level resolvability of cryoEM maps and evaluates the fit between cryoEM maps and their corresponding atomic models. A Q-score of 1.0 represents a perfect fit^27^. Entries with Q-scores below a user-defined threshold are discarded, offering flexible control over map– model quality. In addition, each entry may include one or more cross-references to UniProtKB and AlphaFold, corresponding to functionally annotated protein domains and their structure predictions, respectively^29,30^. To reduce redundancy, CryoDataBot evaluates whether different entries refer to the same protein domain or structure prediction, and employs a two-tier filtering strategy (Methods), consisting of uniqueness filtering and similarity filtering. This strategy removes highly redundant structures while maintaining dataset diversity. By applying these quality control steps, the curated dataset retains only the most informative, high-confidence entries, enhancing data reliability and reducing unnecessary computational burden during downstream processing.

The workflow of the **Structural Data Conditioning** module is illustrated in Figure 1c. Based on the curated metadata, CryoDataBot automatically retrieves structural data files from EMDB and PDB, obtaining, for each entry, a cryoEM map paired with its corresponding atomic model. To facilitate AI training, maps with non-uniform voxel sizes are resampled to user-defined voxel sizes to ensure consistency across the dataset (Methods). After resampling, each map undergoes adaptive density normalization based on its recommended contour level, ensuring consistent density scaling and providing an inherent denoising effect. This method is more flexible and removes more noise than the fixed-threshold approaches commonly used in prior studies (Methods). To further ensure global structural coherence, we did not adopt 3D global correlation scoring methods, which often require intensive voxel-level operations and large memory footprints. Instead, we developed a simple but effective approach: each map–model pair undergoes MMF validation using the Volume Overlap Fraction score (VOF score, see Methods for details), where a high VOF score reflects the accuracy and completeness of the atomic model. Users can apply a configurable threshold to exclude poorly aligned pairs, ensuring that the final dataset consists solely of structurally coherent examples.

Through the **Customized Dataset Construction** module (Fig. 1d), users specify the structures to be labeled, which can include atomic groups (e.g., all atoms within α-helices) or individual atoms (e.g., Cα atoms), along with label values (e.g., 1 or 2). The module generates the corresponding structural labels using the atomic model (Methods). Representative labels are shown in the green panel of Figure 3a and Figure 4. Once the label data is generated, it is paired with the corresponding cryoEM map and partitioned into smaller 3D sub-volumes based on user-defined stride and patch dimensions. These sub-volumes are then split into training, validation, and test sets according to user-defined ratios, creating an AI-compatible dataset for downstream learning applications.

### Construction of Benchmarking Datasets Using CryoDataBot

State-of-the-art automated modeling tools, such as DeepTracer, ModelAngelo, and CryoREAD, leverage deep learning to build atomic models by identifying protein and RNA secondary structures, localizing key backbone atoms, and classifying residue types^12–14^. To systematically evaluate the effectiveness of CryoDataBot in generating high-quality datasets for structure prediction, we constructed three benchmarking datasets with increasing levels of quality control: a **raw dataset** without quality control, a **control dataset** with basic redundancy filtering, and an **experimental dataset** curated through CryoDataBot’s full-quality control pipeline. All three datasets were based on ribosome structures due to their abundance in the EMDB and their inclusion of both protein and RNA components, which are essential for assessing structure prediction across multiple molecule types.

A total of 962 ribosome-related entries were initially retrieved from EMDB using the query “ribosome AND resolution:[3 TO 4]”. Following the exclusion of 20 entries due to unsuccessful normalization, 942 entries remained. This unfiltered collection served as the raw dataset, mimicking prior methodologies such as Cryo2StructData^25^. It also provided the foundational pool from which the control and experimental datasets were derived through successive quality control steps. Detailed metadata and a summary of the filtration process are provided in the Supplementary Tables.

Table 2 summarizes the number of entries discarded at each quality control stage for both the control and experimental datasets. For the control dataset, a uniqueness filter was applied to remove 173 fully redundant entries that shared identical UniProtKB cross-references, yielding 769 entries, from which 18 were randomly selected as a holdout test subset (these entries are not included in the experimental dataset), yielding a final control dataset of 751 entries.

**Table 2:** Quality control (QC) stages applied to construct the raw, control and experimental datasets. Entry counts are shown before and after QC stages, with values in parentheses indicating the number of entries discarded during each stage. After undergoing the full QC pipeline, the final experimental dataset comprises only 143 entries. This was achieved by eliminating highly redundant and structurally incompatible entries, thereby yielding a more compact, reliable dataset for training.

| Entry Count (Pre-QC) | 942 |  |  |
| --- | --- | --- | --- |
| Quality Control Stages | Raw dataset | Control dataset | Experimental dataset |
| Q-score Filter (Threshold: 0.4) | X | X | ✓ (-406) |
| Uniqueness Filtering | X | ✓ (-173) | ✓ (-92) |
| Similarity Filtering (Threshold: 70%) | X | X | ✓ (-230) |
| MMF Validation (Threshold: 0.82) | X | X | ✓ (-71) |
| Entry Count (Post-QC) | 942 | 769 | 143 |

The experimental dataset underwent a more rigorous, multi-stage quality control pipeline (Table 2). First, entries with Q-scores below 0.4 were discarded, removing 406 low-quality entries. In the first tier of redundancy filtering, 92 entries were discarded: 23 due to invalid cross-references and 69 due to duplication identified through identical UniProtKB annotations. In the second tier, an additional 230 entries exhibiting over 70% similarity were discarded based on UniProtKB-derived similarity evaluation. These quality control steps yielded an experimental dataset of 214 high-confidence, low-redundant entries—approximately one-fourth the size of the raw and control datasets—thus reducing computational overhead while preserving data informativeness. Figure 1b presents representative examples of excluded entries, highlighting improvements in data quality through the removal of structurally incoherent and redundant entries.

Structural data—including cryoEM maps and atomic models—were collected from EMDB and PDB for all three datasets. All cryoEM maps were then resampled to a voxel size of 1 Å, denoised according to their recommended contour levels, and normalized to a 0–1 density scale to ensure uniformity. For the experimental dataset, CryoDataBot further assessed the MMF using the VOF score, excluding an additional 71 entries that fell below the 0.82 threshold (Table 2, Fig. S1). This refinement yielded a final set of 143 high-fidelity map–model pairs for the experimental dataset. In comparison, the preprocessed control and raw datasets contained 751 and 942 map–model pairs, respectively.

Using the preprocessed structural data, secondary structure labels were generated for both the control and experimental datasets. The labeled volumes, together with the normalized cryoEM maps, were partitioned into 64^3^Å^3^ sub-volumes to support batch training. These sub-volumes were subsequently split into training and validation sets at an 80:20 ratio. One entry from the control dataset (EMD-2875) was excluded due to inconsistencies between its cryoEM map and label data, rendering it unsuitable for model training. The resulting control and experimental datasets were thus fully prepared for direct use in U-Net model training workflows.

### Quality Assessment of the Constructed Benchmark Datasets

High-quality training datasets are essential for deep learning-based structural modeling. Two critical attributes that determine dataset utility are map–model fitness (MMF) and structural redundancy. Accurate map–model alignment ensures reliable label generation from atomic models, while low redundancy enhances dataset diversity and reduces overfitting risks during model training. To evaluate these attributes, we systematically assessed the raw, control, and experimental datasets curated in this study.

To evaluate MMF, we computed multiple correlation coefficient (CC) metrics for each map–model pair using PHENIX^9^, including CC_mask (for atomic center fit), CC_volume (for molecular envelope fit), CC_peaks (for fit of strong peaks), and CC_box (for overall map similarity). These metrics reflect distinct aspects of map–model consistency. As shown in Figure 2a, the experimental dataset consistently outperformed both the raw and control datasets across all CC metrics. Both the raw and control datasets contained a significant number of map–model pairs with CC values below 0.6, indicating poor correspondence between cryoEM maps and atomic models. In contrast, the experimental dataset exhibited fewer low-CC entries and consistently higher 25th, 50th, and 75th percentile values across all CC metrics. These results highlight the effectiveness of Q-score filtering and MMF validation in improving map–model consistency within the dataset, thereby enabling the generation of precise structural labels.

**Fig. 2:**
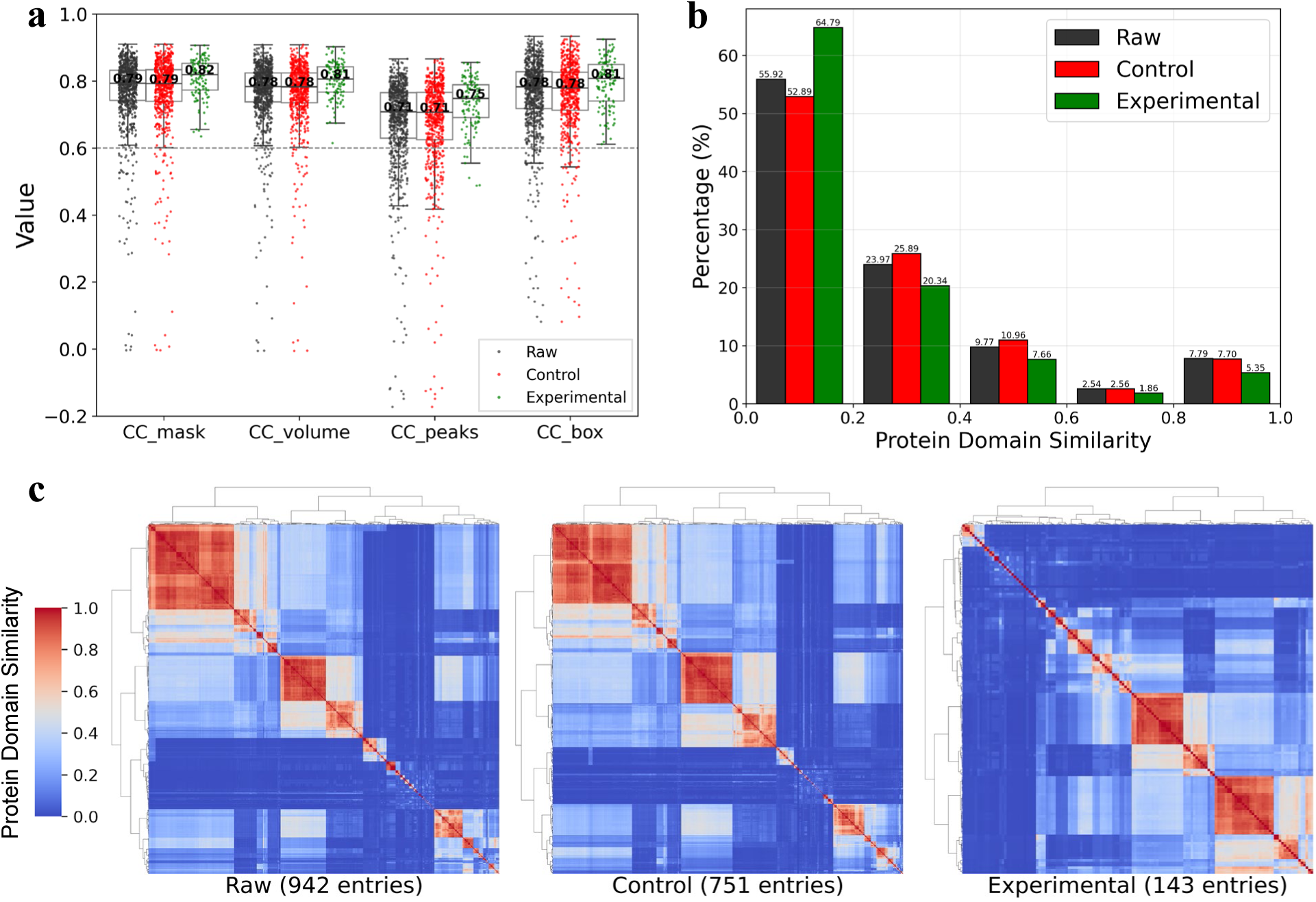
Evaluation of map–model fitness and structural redundancy in raw, control, and experimental datasets. **a**, Correlation coefficient (CC) metrics showing that the experimental dataset (green) consistently outperformed both the raw (dark gray) and control (red) datasets. The statistics were calculated over n = 942, 751, and 143 independent cryoEM maps in the raw, control, and experimental datasets, respectively. The numbers in the middle of the box plots represent the median values for each CC metric across the datasets. The gray dashed line in the figure represents a CC value of 0.6. **b**, A histogram illustrating the percentage distribution of pairwise protein domain similarity scores across the three datasets. The x-axis indicates similarity score ranges, with the values above each bar denoting the proportion of each dataset within those ranges. The experimental dataset shows the highest proportion of least similar pairs (below 0.2) and the lowest proportions of mid- (0.2-0.6) and high-similarity pairs (above 0.8), indicating reduced redundancy. **c,** Clustered heatmaps of the pairwise protein domain similarity matrix across the three datasets, with warmer colors indicating higher similarity. The raw and control datasets show dense clusters of high-similarity pairs (values > 0.5), while the experimental dataset shows a more dispersed and localized distribution of high-similarity pairs.

To assess dataset redundancy, we analyzed structural similarity between entries based on InterPro (IPR) domain annotations^31^. IPR identifiers, which represent conserved protein domains with independent folding and functional capabilities, were retrieved for each entry using its PDB ID. Pairwise similarity scores were computed as the ratio of shared IPR identifiers between entry pairs. Figure 2b displays the distribution of similarity scores across all datasets. The experimental dataset contains the highest proportion (64.79%) of low-similarity pairs (similarity score < 0.2), and the lowest proportion of highly similar pairs (similarity score > 0.6), indicating the lowest overall redundancy. This pattern is further supported by the similarity heatmaps in Figure 2c, where both the raw and control datasets exhibit dense clusters of high-similarity pairs (scores > 0.5), reflecting substantial redundancy from entries with partial or complete structural similarity. In contrast, the experimental dataset shows a more dispersed and localized distribution of high-similarity scores, demonstrating the effectiveness of the redundancy-filtering strategy. To assess redundancy from a complementary perspective, we defined sequence-based similarity between entries using the result from BLASTp^32^ (see Fig. S2 for details). The results (Fig. S2), consistent with our earlier structural similarity analysis, confirm that the filtering strategy effectively reduces redundancy at the sequence level as well.

A comparison between the raw and control datasets provides additional insight. The control dataset contains a lower proportion of entry pairs with similarity scores in the 0.8–1.0 range (Fig. 2b), demonstrating the effectiveness of the uniqueness filtering stage in eliminating exact duplicates. However, it has a higher proportion of mid-similarity pairs (0.2–0.6) and a lower proportion of low-similarity pairs (below 0.2). The increase in mid-similarity proportions likely arises from the preservation of partially similar entries, combined with a decrease in the overall number of entry pairs. Moreover, the excluded redundant entries could have formed low-similarity pairs when matched with structurally unrelated entries; their removal reduces such combinations, contributing to the observed decline in low-similarity proportions. These results suggest that although uniqueness filtering effectively removes exact duplicates, it does not fully address broader structural similarities.

### Performance of U-Net Trained on CryoDataBot-Generated Datasets

To directly assess how dataset quality influences model training and predictive performance, we trained two identical 19-layer 3D U-Net models on the control and experimental datasets, respectively. These models, commonly utilized in AI-based structural modeling frameworks^19^, were designed in this study to predict secondary structures from cryoEM maps (Fig. 3a). Both U-Net models were trained under identical conditions with consistent learning rates and early stopping criteria. Throughout training and upon completion, model performance was systematically assessed across multiple evaluation dimensions (Methods).

**Fig. 3:**
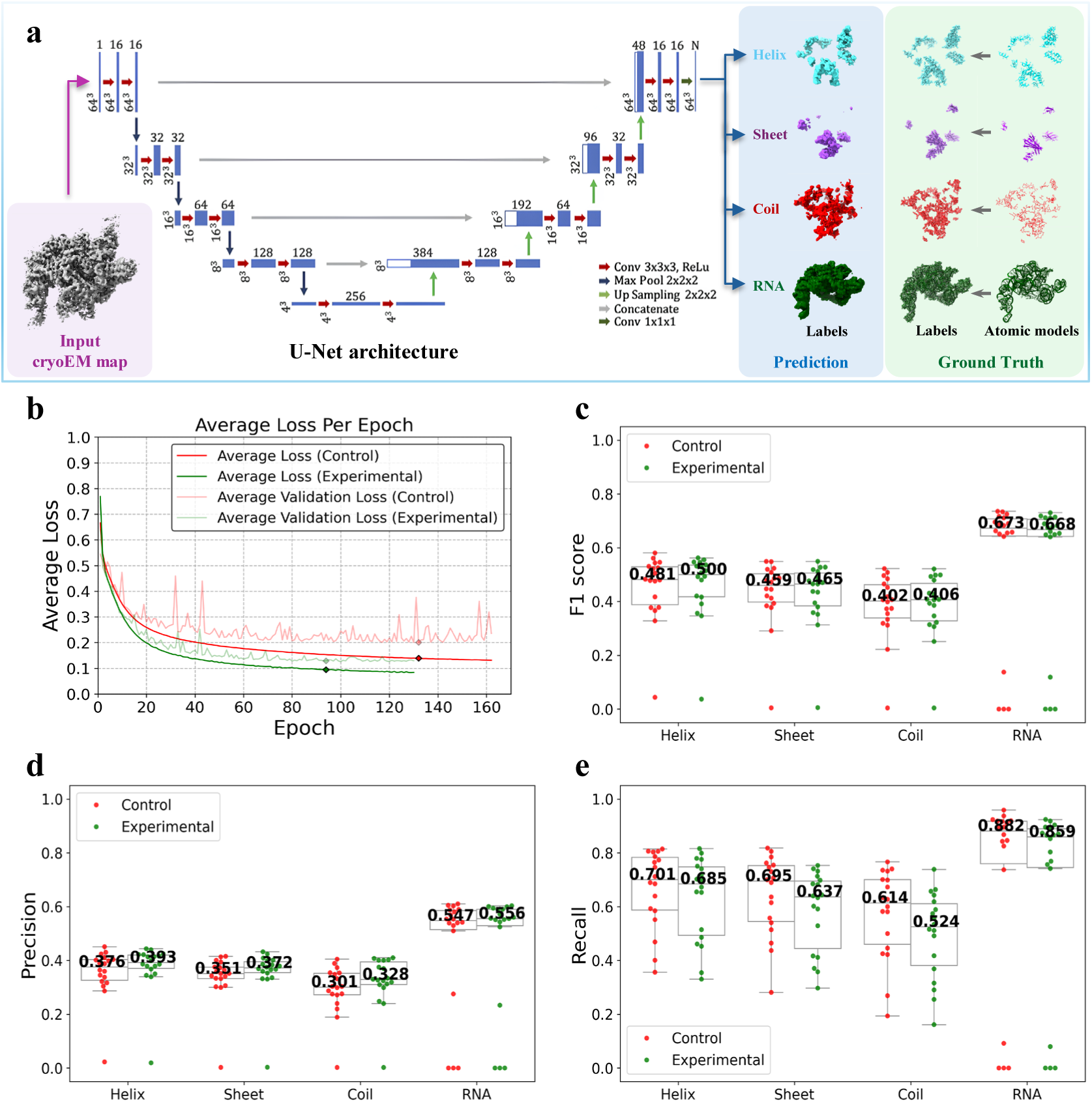
Evaluation of dataset performance in U-Net model training. **a,** Schematic of a 19-layer 3D U-Net architecture for secondary structure prediction from cryo-EM maps. The input (purple panel) is the cryoEM map, and the output (blue panel) contains predicted labels for helices, sheets, coils, and RNA. Ground truth labels derived from atomic models are shown in the green panel. The central diagram illustrates the U-Net architecture, adapted from DeepTracer. Blue bars represent feature maps, with spatial dimensions indicated at the bottom left and the number of channels at the top. **b**, Average loss curves for control (red) and experimental (green) training. Solid lines represent training loss, and lighter lines represent validation loss. The control and experimental models were trained for 162 and 130 epochs, respectively, with the best-performing epochs marked at 132 and 94 (diamonds). **c**, F1 scores for each structural label, calculated on an independent test set of n = 18 cryoEM maps using the U-Net model trained on the control dataset (red) and the experimental dataset (green) at their respective best-performing epochs. The numbers at the middle of the box plots indicate the median values. **d-e**, Same as **c**, but for precision and recall, respectively.

As detailed in Table 3, the experimental dataset outperformed the control dataset in improving training efficiency. Models trained on the experimental dataset achieved faster convergence and higher learning efficiency, requiring significantly fewer epochs (130 vs. 162) to reach early stopping. Figure 3b corroborates this accelerated convergence, showing a steeper and more consistent decline in loss per epoch for the experimental training (green line). In addition to faster convergence, the experimental training also demanded significantly less computation, with each epoch completing in just 30 minutes versus 87 minutes for the control—a reduction attributed to the dataset’s smaller size. Collectively, these improvements resulted in a more than threefold decrease in total training time (2.7 vs. 9.8 days), substantially accelerating the overall model development process.

**Table 3.** Comparison of U-Net model training efficiency and post-training performance on the control and experimental datasets. Early stopping epochs, training time per epoch, and total training time were systematically collected to evaluate training efficiency. Post-training evaluation was performed on the validation set using the best-performing epochs (epoch 132 for the control dataset and epoch 94 for the experimental). Key performance metrics, including overall accuracy, precision, recall, and F1 score, were calculated to assess model prediction performance. F1 scores for helix, sheet, coil, and RNA predictions appear in parentheses. Across all evaluation criteria, the experimental dataset consistently demonstrated superior performance over the control dataset.

| Training set | Control dataset | Experimental dataset |
| --- | --- | --- |
| Early stopping epochs | 162 | 130 |
| Training time per epoch (min) | 87 | 30 |
| Total training time (day) | 9.8 | 2.7 |
| Accuracy | 88.06% | 90.14% |
| Precision | 0.41 | 0.46 |
| Recall | 0.96 | 0.98 |
| F1 score | 0.57 | 0.62 |
| F1 score for each label | (0.51, 0.54, 0.47, 0.63) | (0.58, 0.60, 0.57, 0.66) |

At the best-performing epochs (epoch 132 for the control training and epoch 94 for the experimental), the overall accuracy, precision, recall, and F1 score (Methods), all calculated on the validation set, are summarized in Table 3. Despite its substantially smaller size, the experimental dataset enabled the model to outperform the control across all evaluated metrics, achieving higher accuracy (90.14% vs. 88.06%), precision (0.46 vs. 0.41), recall (0.98 vs. 0.96), and F1 score (0.62 vs. 0.57), with consistently better F1 scores across all structural labels. The better performance of the model trained on the experimental dataset was consistently evident throughout training (Fig. S3).

The previously defined independent test set of 18 cryoEM maps and their corresponding atomic models, randomly selected before training and excluded from both datasets, was used to evaluate both U-Net models. Precision, recall, and F1 scores were independently computed for each structural label within each map (Fig. 3c-e). While the overall F1 scores indicated comparable performance between the experimental and control models, the experimental model consistently demonstrated higher precision across all structural labels, reflecting more accurate and reliable predictions. The slightly lower recall of the experimental model reflects the precision–recall tradeoff. The lower recall of the experimental model reflects a slight decrease in true positives (Fig. S4), which may result from the reduced sample size and sample diversity introduced by stringent filtering. In contrast, although the number of true positives decreases, the stronger effect of reduced false positives due to cleaner and more conservative training data (Fig. S4) leads to higher precision and an overall high F1 score. Considering this tradeoff, users should calibrate decision thresholds to their use case such as relaxing thresholds when recall is paramount. Thresholds may vary depending on the type and quality of the target data, so users are advised to grid search quality filter settings to determine which thresholds best fit their purpose. For example, a de novo modeling software dependent on capturing the entire structure might warrant higher recall performance. Overall, the improved precision suggests a favorable tradeoff, resulting in a model that is more robust and less prone to generating false positives.

Of the 18 test set examples, Figure 3c-e shows that sometimes, prediction results may have poor precision and recall, quantified by the outlier dots lying outside the interquartile range of the box plot. Figure 4 presents two representative test-set examples explaining this observation for the experimental U-Net model: EMD-32074, with the lowest recall, and EMD-3245, with the lowest precision. Blue panels show the predicted secondary structures, restricted to regions with prediction probabilities > 0.8. Green panels display the corresponding atomic models—manually built by the original authors and deposited in the PDB—along with their derived labels, treated as “ground truth”. Evaluation metrics discussed earlier reflect the agreement between these predicted and “ground truth” labels. In the low-recall case (EMD-32074, PDB ID: 7VPX), the deposited atomic model contains substantial inaccuracies, with numerous atoms placed in regions lacking significant cryoEM density. As a result, the “ground truth” labels only partially reflect the actual map features. In contrast, the U-Net model confines its predictions to high-density regions, correctly avoiding unsupported areas, suggesting that the low recall arises from ground truth errors rather than model limitations. Conversely, in the low-precision case (EMD-3245, PDB ID: 3JC2), the atomic model covers only a limited portion of the map, leaving extensive regions of cryoEM density unlabeled. Nevertheless, the U-Net model correctly identifies structural features across the map, such as continuous, well-resolved RNA helices. The resulting low precision thus reflects incomplete “ground truth” labels rather than inaccurate predictions. Together, these examples highlight the U-Net model’s robustness against incomplete or noisy ground truth and its potential utility in refining or validating deposited atomic models by directly leveraging cryoEM maps. Notably, these examples—cryoEM map and model pairs—are associated with very low Q-scores or VOF scores, which suggests an important implication: CryoDataBot can function not only as a data preparation pipeline but also as a potential quality control tool. When entries are excluded during the Q-score filtering or MMF validation stage, it may indicate that the corresponding atomic model does not fit well with the cryoEM map and may warrant further manual inspection.

**Fig. 4:**
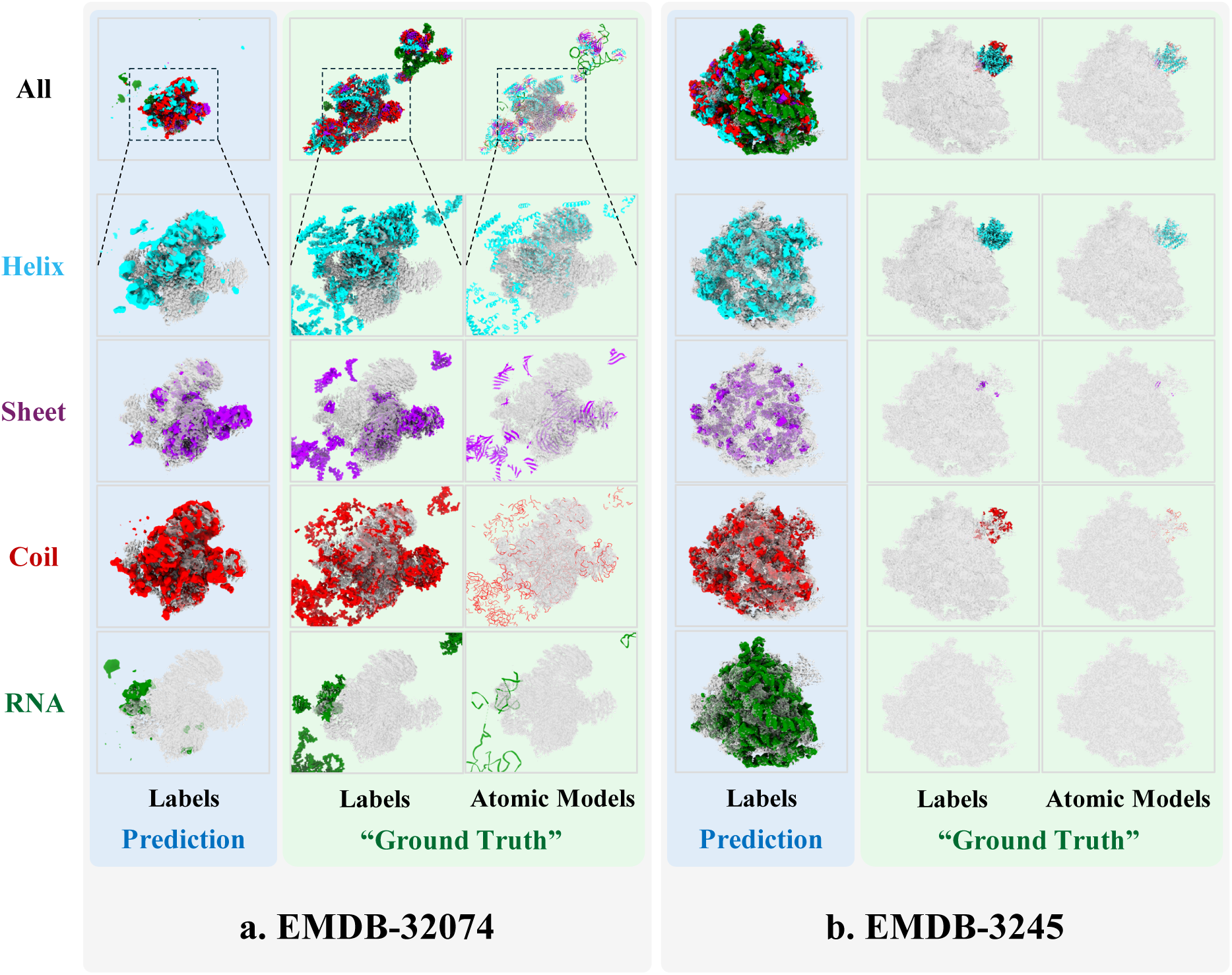
Representative predictions from the U-net model trained on the experimental dataset, showing one example with low recall and another with low precision on the test set. Predicted labels—including composite outputs and separate annotations for helix, sheet, coil, and RNA—are shown alongside the corresponding “ground truth”, derived from manually built atomic models in the PDB. Blue panels show predictions; green panels show “ground truth”. CryoEM maps are rendered as transparent overlays. Color code: cyan (helix), purple (sheet), red (coil), green (RNA). **a,** Example from EMDB-32074, where substantial portions of the ground truth atomic model are built without supporting cryoEM map density. The U-net model correctly abstains from predicting in these areas, leading to low recall. **b**, Example from EMDB-3245, where large regions of the cryoEM map remain unmodeled in the ground truth. The U-net model predicts plausible structures in these areas, resulting in low precision when evaluated against the incomplete ground truth.

### Practical Validation: Retraining CryoREAD with CryoDataBot-Generated Datasets

To further demonstrate the applicability of CryoDataBot-generated datasets within established modeling frameworks, we retrained Stage 1 of CryoREAD—a deep learning framework for de novo modeling of DNA and RNA atomic structures from cryoEM maps^14^. As shown in Figure 5a, Stage 1 is designed to identify and classify key atomic groups—namely sugar, phosphate, base, and base types (A, U/T, C, G)—from the input cryoEM map. In the original CryoREAD study, the Stage 1 training set was constructed through a complex and labor-intensive pipeline involving EMDB map collection, redundancy reduction via clustering, voxel resampling to 1.0 Å, density normalization, and sub-volume extraction, ultimately comprising 290 distinct RNA maps^14^. To replicate this format, we used CryoDataBot to automatically generate new structural labels based on our experimental ribosome dataset consisting of 143 maps, as described in the previous section. The entire process was fully automated by configuring a small set of parameters—including atomic group selection (sugar, phosphate, A, U/T, C, G), resampling voxel size (1.0 Å), labeling radius (2.0 Å), and sub-volume size (64³ Å³)—without requiring further manual intervention. Training was performed using the same hyperparameter settings as in the original CryoREAD training. Evaluation was conducted on a consistent test set comprising 63 entries, obtained by removing five overlapping or invalid cases (EMD-10535, EMD-6789, EMD-21856, EMD-9572, and EMD-4138) from the original 68.

**Fig. 5:**
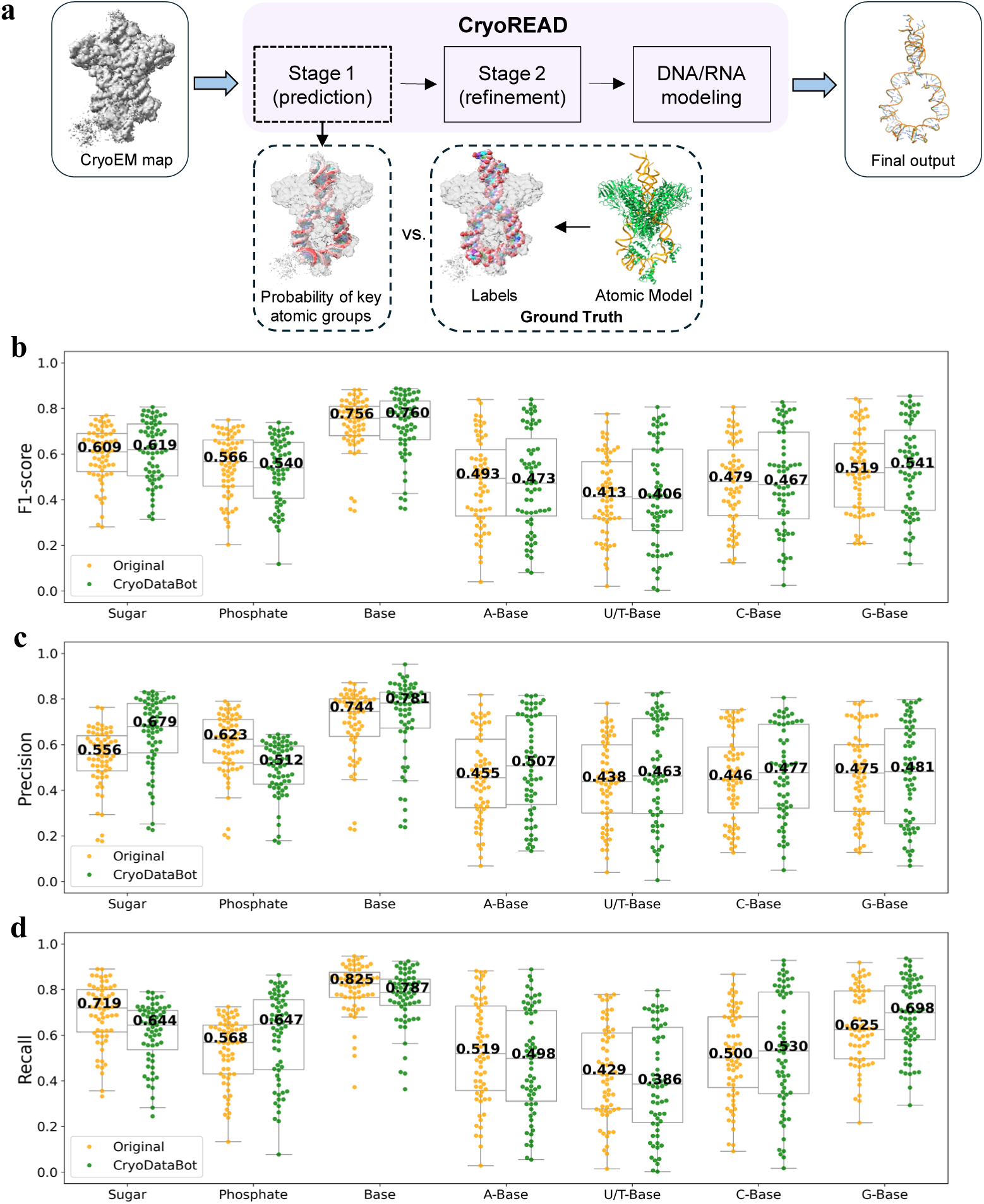
Comparative evaluation of prediction performance between the original and retrained CryoREAD Stage 1 models. **a**, Overview of the CryoREAD workflow. The input is a cryoEM map and the final output is an automatically built DNA or RNA atomic model. Stage 1 of CryoREAD identifies and classifies key atomic groups (sugar, phosphate, base, and base types), and its predictions are compared against reference labels derived from the deposited atomic model (example entry: EMD-7480, PDB ID: 6CIJ). **b,** F1 scores for each structural label, calculated on an independent test set of n = 63 cryoEM maps. Results are shown for the original CryoREAD model (yellow) and the retrained version using the CryoDataBot-generated dataset (green). Median values are indicated within each box plot. **c-d**, Same as **b**, but for precision (**c**) and recall (**d**) metrics.

Evaluation results for the retrained Stage 1 model are shown in Figures 5b-d (green), alongside those of the original model (yellow). For structure detection, the median F1 scores (Fig. 5b) for sugar, phosphate, and base are 0.619, 0.540, and 0.760, respectively, reflecting the model’s ability to accurately localize fundamental components of nucleic acid structures. For fine-grained base classification, the median F1 scores for individual base types—A, U/T, C, and G—are 0.473, 0.406, 0.467, and 0.541, respectively. These results indicate that the retrained model achieves approximately 50% accuracy for each base type, substantially exceeding the 25% expected by random chance among four classes. These scores are comparable to those of the original model, demonstrating that CryoREAD retains high predictive performance when trained on CryoDataBot-generated datasets, despite the reduced dataset size, absence of manual curation, and restriction to a single sample type—ribosome-derived structures.

The precision (Fig. 5c) and recall (Fig. 5d) for structure detection again reveal a tradeoff. Compared to the original training, training on CryoDataBot-generated dataset results in higher precision but lower recall for detecting sugar (precision: 0.679 vs. 0.556; recall: 0.644 vs. 0.719) and base (precision: 0.781 vs. 0.744; recall: 0.787 vs. 0.825). However, phosphate detection shows the opposite trend, with lower precision (0.512 vs. 0.623) but higher recall (0.647 vs. 0.568) when trained on CryoDataBot-generated dataset. The observed changes in precision and recall are non-negligible, underscoring that training set composition can differentially affect model performance. This highlights the importance of selecting training data carefully according to study-specific goals, with attention to factors such as structural diversity, data quality, and preprocessing settings.

Figure 6 illustrates a representative example (EMD-3532) of RNA atomic group prediction by the retrained CryoREAD Stage 1 model. As in previous evaluations, the ground truth was derived from the deposited atomic model, manually built by the original authors and available in the PDB. The green panel shows the cryoEM map (gray) overlaid with the ground truth model (spheres). The blue panels (Fig. 6b–g) present Stage 1 predictions, rendered as transparent volumes with probability > 0.4, overlaid with the ground truth atomic model in ball-and-stick format to facilitate direct structural comparison. The composite prediction (Fig. 6a) omits regions corresponding to protein density—as indicated by the white cartoon representation of the protein model in Figure 6h—and accurately reconstructs the canonical RNA double helix. The sugar (pink) and phosphate (red) groups form the backbone, while the four types of bases are embedded between the helices. The predicted structure closely aligns with the ground truth, not only in the overall RNA topology but also in the precise localization of atomic groups and accurate classification of base types (Fig. 6b-g). This high level of accuracy provides a strong foundation for downstream automated modeling stages in CryoREAD. Overall, the example demonstrates the effectiveness of deep learning in interpreting complex cryoEM maps and underscores the robustness of the CryoDataBot-generated training data in enabling reliable structure prediction.

**Fig. 6:**
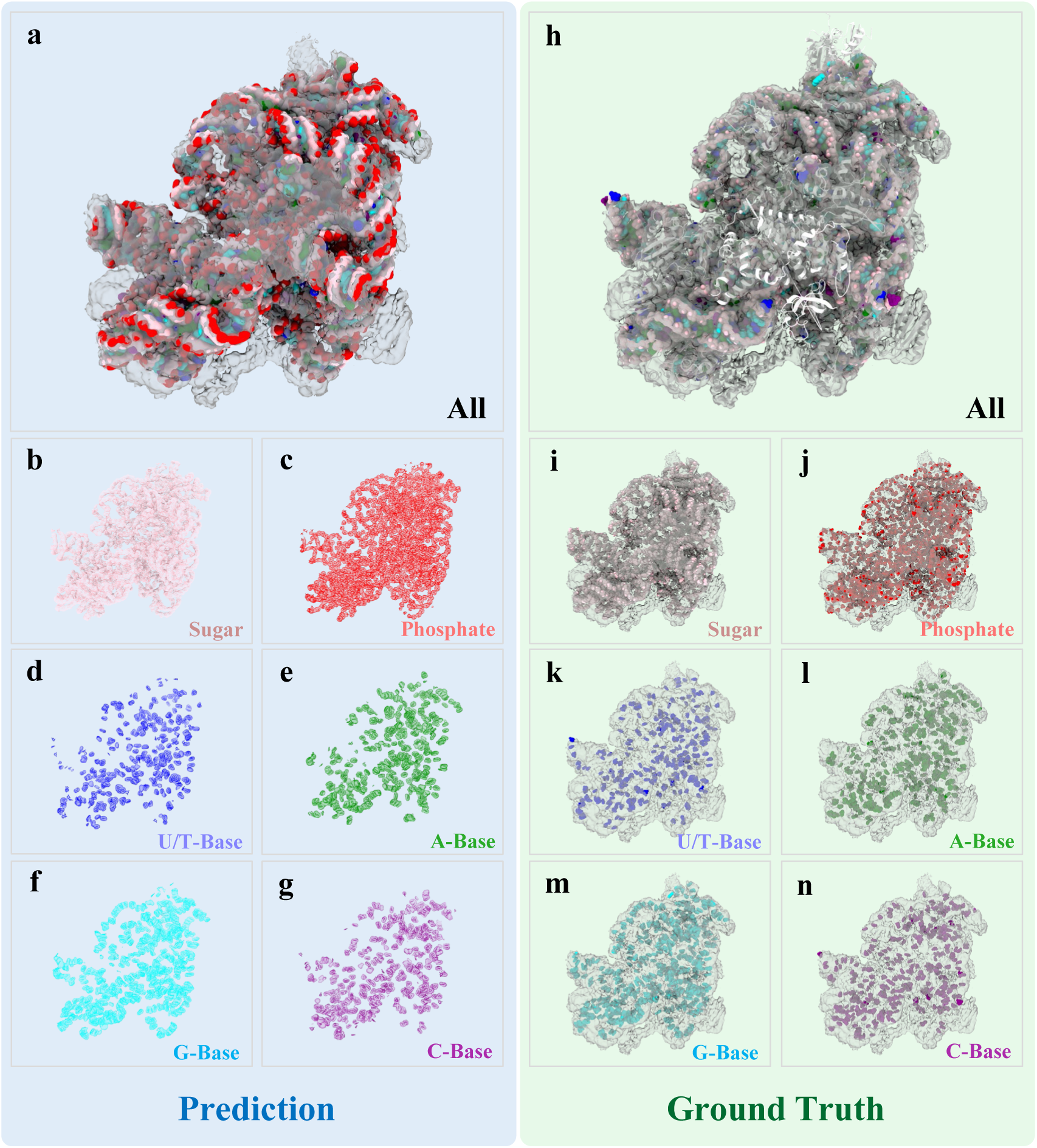
Representative example (EMD-3532) of a test-set prediction from the retrained CryoREAD model. CryoREAD predictions (regions with probability > 0.4) are shown in the blue panels, while the corresponding ground truth atomic models are shown in the green panels. Different colors indicate different atomic groups: pink for sugar, red for phosphate, blue for U/T, green for A, light blue for G, purple for C, and white for protein (shown only in panel **h**). **a**, Composite prediction (colored volumes) overlaid with the original cryoEM map (gray). **b–g**, Individual predictions (transparent volumes) for each atomic group, overlaid with the ground truth atomic model (ball-and-stick representation). **h**, Ground truth atomic model, with RNA shown as spheres and protein as a cartoon, overlaid with the cryoEM map (gray). **i-n,** Ground truth representations for individual atomic groups (shown as spheres), overlaid with the original cryoEM map (gray).

Further comparison reveals that the restriction to a single sample type—ribosome-derived structures—has two notable effects. First, it reduces generalizability: as shown in Figure S5, the retrained model performs consistently better on ribosome cases (38 out of 63 test maps) than on other RNA/DNA classes (25 maps), whereas the original CryoREAD model (Fig. S6), trained on a more diverse set of RNA/DNA structures, shows a smaller performance gap between ribosome and non-ribosome cases. Second, it enhances CryoREAD’s predictive performance specifically for ribosome structures, underscoring the importance of tailoring training data to the target application in structure-specific studies.

### Robustness Check: Benchmarking on G protein Dataset

To demonstrate the broad applicability of CryoDataBot, we generated datasets based on G proteins—key regulators of cellular signal transduction that are notably smaller in size compared to ribosomes. Using the search keywords “G Protein-Coupled Receptors AND resolution:[3 TO 4}”, we retrieved 152 metadata entries from EMDB (Table 4). From these, a cluster of 26 similar entries was reserved as the test set, while the remaining 126 entries constituted the raw dataset. We applied the same metadata curation thresholds used for ribosomes (Q-score ≥ 0.4, similarity ≤ 70%) and adopted identical structural conditioning settings. However, to ensure a sufficient number of entries for training—given the smaller size of the G protein dataset relative to the ribosome dataset (942 entries)—we employed a more lenient MMF filter of 0.65, resulting in an experimental (Q0.4) dataset comprising 38 entries. Additionally, we constructed two comparative experimental groups using Q-score thresholds of 0.3 and 0.5, yielding the Q0.3 and Q0.5 datasets with 48 and 16 entries, respectively (Table 4). For the purpose of training U-Net models to identify C-alpha positions, C-alpha (“CA”) was chosen as the target for labeling. Using the same settings as for ribosomes, we performed partitioning and data splitting to generate both training and validation sets for the raw, experimental, Q0.3 and Q0.5 datasets. U-Net models were trained using the same settings as used for ribosome except for the batch number and output classes (Methods).

**Table 4:**
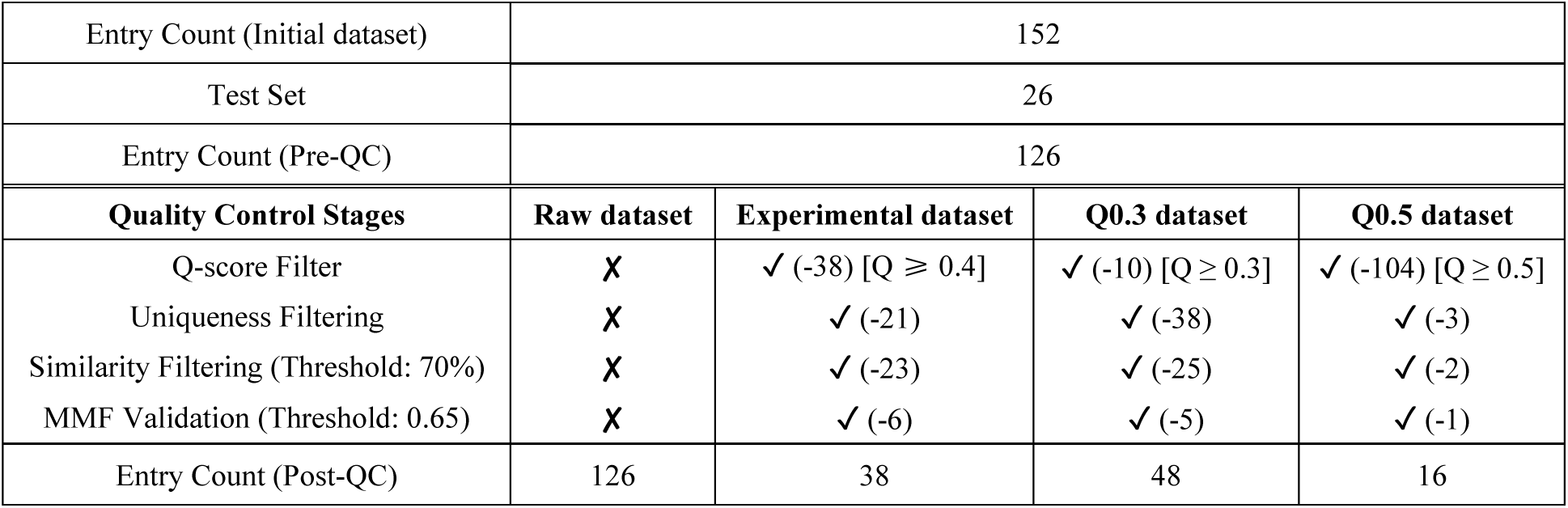
Similar to Table 2, but for constructing datasets with G protein data. A randomly generated test set was first removed from the initial 152 entries. The threshold of MMF validation was set to 0.65 to retain more entries in the experimental training set.

| Entry Count (Initial dataset) | 152 |  |  |  |
| --- | --- | --- | --- | --- |
| Test Set | 26 |  |  |  |
| Entry Count (Pre-QC) | 126 |  |  |  |
| Quality Control Stages | Raw dataset | Experimental dataset | Q0.3 dataset | Q0.5 dataset |
| Q-score Filter | ✗ | ✓ (-38) [$Q \geq 0.4$] | ✓ (-10) [$Q \geq 0.3$] | ✓ (-104) [$Q \geq 0.5$] |
| Uniqueness Filtering | ✗ | ✓ (-21) | ✓ (-38) | ✓ (-3) |
| Similarity Filtering (Threshold: 70%) | ✗ | ✓ (-23) | ✓ (-25) | ✓ (-2) |
| MMF Validation (Threshold: 0.65) | ✗ | ✓ (-6) | ✓ (-5) | ✓ (-1) |
| Entry Count (Post-QC) | 126 | 38 | 48 | 16 |

Compared to training with the raw dataset, the other three U-Net models converged more rapidly, exhibiting a steeper and more consistent decline in loss per epoch, and reaching early stopping in nearly half the number of epochs (Fig. 7a). In the Q0.3, experimental (Q0.4), and Q0.5 datasets, CryoDataBot filters out low-consistency cryoEM map–model pairs, resulting in smaller datasets with more precise labels that enhance training efficiency. Notably, higher Q-score thresholds lead to smaller datasets and faster convergence. This trend is also reflected in the training dynamics (Fig. S7), where models trained on higher Q-score datasets achieve higher overall precision and F1 scores in fewer epochs, driven by improved label accuracy in both the training and validation sets under stricter filtering.

**Fig. 7:**
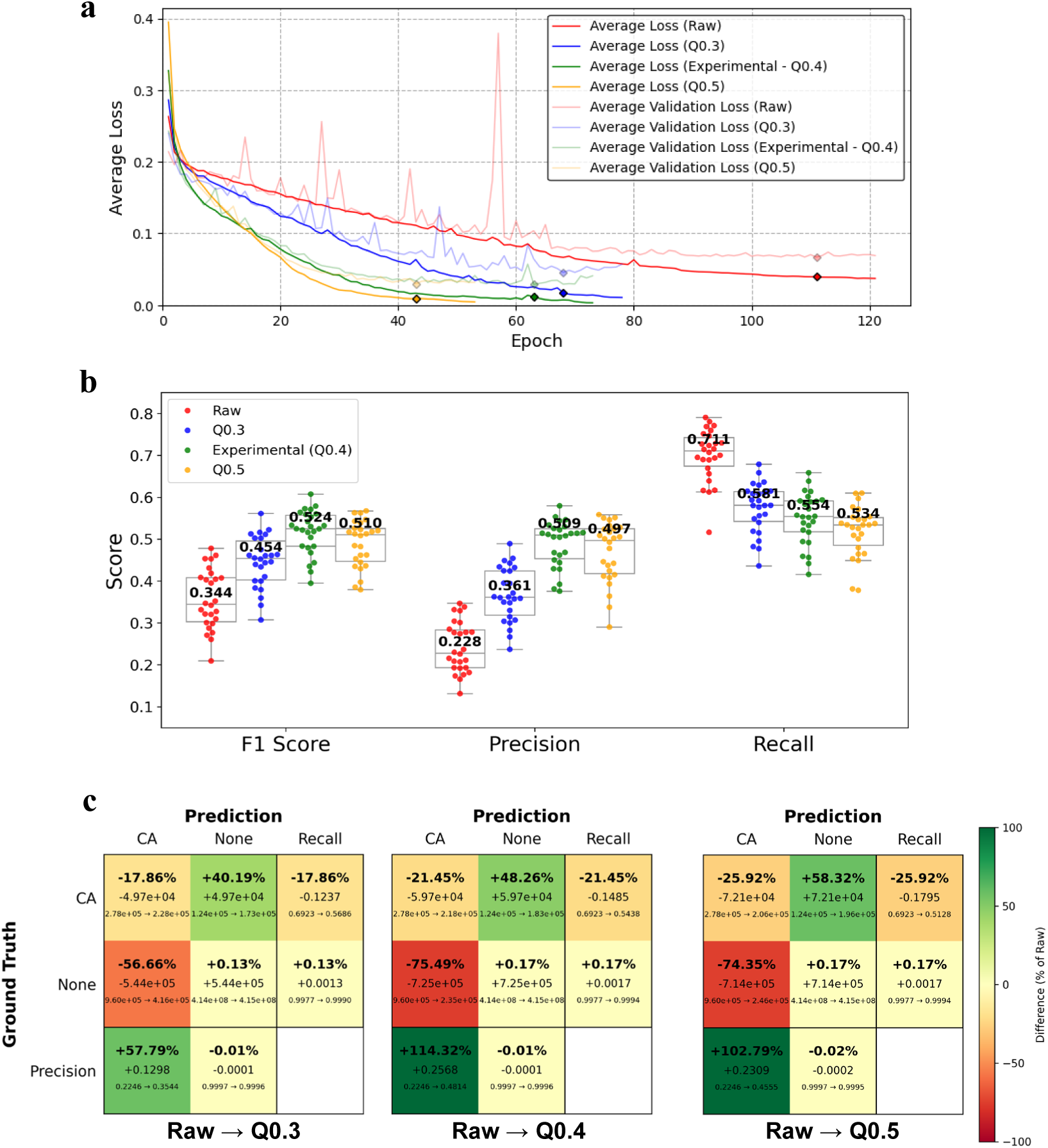
Evaluation of G protein dataset performance in U-Net model training. **a,** Average loss curves for models trained on four datasets: raw (red), Q0.3 (blue), experimental (green), and Q0.5 (orange). Solid lines represent training loss, and lighter lines represent validation loss. The models were trained for 121, 78, 73, and 53 epochs, respectively, with their best-performing epochs marked by diamonds at 111, 68, 63, and 43. **b**, F1 score, precision, and recall evaluated on an independent test set of *n* = 26 cryoEM maps, using U-Net models trained on the four datasets at their respective best-performing epochs. Median values are shown at the center of each box plot. **c**, Confusion matrix comparisons from the raw model to Q0.3, experimental (Q0.4), and Q0.5 models, shown from left to right. Each cell contains three lines: (bottom) raw model count → current model count, (middle) absolute difference (current minus raw), and (top) ratio of difference to raw. Cells are color-coded based on this ratio. True positive counts of CA were slightly reduced by 17.86%, 21.45% and 25.92%, and false positive counts largely reduced by 56.66%, 75.49%, and 74.35% in Q0.3, Q0.4 and Q0.5 training models, respectively.

F1 score, precision, and recall evaluated on an independent test set of *n* = 26 cryoEM maps are shown in Figure 7b. Models trained on the Q0.3, Q0.4, and Q0.5 datasets consistently outperformed the raw-data model in terms of F1 score, underscoring the robustness of CryoDataBot in enhancing data quality. A familiar precision–recall tradeoff emerges: the raw model achieves the highest recall but suffers from the lowest precision among all models. Confusion matrices in Figure 7c further illustrate this trend. As the dataset becomes more refined, the number of true positives decreases slightly—by 17.86%, 21.45%, and 25.92% for Q0.3, Q0.4, and Q0.5, respectively—resulting in a modest reduction in recall (∼20%). However, the drop in false positives is far more substantial—56.66%, 75.49%, and 74.35%, respectively— leading to marked improvements in precision (∼100%) and overall F1 score.

A comparison across different Q-score thresholds reveals a clear trend: as the Q-score filter becomes more stringent, model recall decreases. This is likely due to reduced dataset size, which in turn limits structural diversity. Interestingly, precision peaks at the moderate threshold of Q-score = 0.4, and the F1 score also reaches its highest value at this intermediate setting. Based on these findings, we recommend using the default Q-score filter of 0.4 when constructing datasets with CryoDataBot, followed by a grid search over alternative thresholds to optimize performance for the specific task and data characteristics.

## Discussion

To meet the critical need for high-quality cryoEM datasets in AI-driven structural modeling, we developed CryoDataBot, which, to our knowledge, is the first software specifically designed to streamline and scale the generation of standardized, high-quality datasets essential for robust model development. Many widely adopted AI frameworks still rely on datasets that are either unfiltered or lack systematic quality control, introducing biases and limiting the ability to fairly evaluate and compare model architectures across tools. CryoDataBot addresses this challenge by automating the dataset construction and enforcing rigorous, standardized quality control. This not only improves the reliability of downstream AI models but also enables fair benchmarking based on a consistent dataset foundation. Furthermore, CryoDataBot offers extensive flexibility, allowing users to configure nearly all processing parameters with fine granularity based on specific modeling objectives and available data, thereby accommodating the diverse needs of structural biology tasks.

We demonstrate that AI models trained on datasets generated by CryoDataBot achieve notably higher precision, reflecting more accurate and reliable predictions. However, models trained on strictly filtered datasets show a modest decrease in recall. This tradeoff likely stems from the reduced dataset size resulting from rigorous filtering, which enhances accuracy but limits exposure to structural variability. Researchers are therefore encouraged to adjust the quality control stringency according to their scientific objectives. For applications requiring high-confidence predictions—such as identifying accurate protein sequences from unknown cryoEM maps—a rigorously filtered dataset is preferable. In contrast, when the goal is to capture broader structural features, such as in automated de novo modeling where further refinement is expected, a more relaxed threshold may improve recall by preserving diversity.

As AI continues to transform cryoEM-based structural biology, the ability to automatically generate high-quality, customizable datasets is becoming essential for next-generation AI tool development. CryoDataBot fulfills this need as a dedicated solution that reduces the technical burden of dataset construction and facilitates broader integration of AI into structural biology research.

The introduction of CryoDataBot marks a significant advancement in cryoEM data handling, offering a robust tool that standardizes dataset creation and supports the growing intersection of structural and computational biology. By ensuring high-quality, curated data, CryoDataBot helps overcome one of the key limitations in AI-based cryoEM modeling: dependence on unreliable or incomplete data. This work not only streamlines dataset preparation but also lays the foundation for more precise and reproducible model predictions. As AI tools evolve, CryoDataBot’s flexibility will be critical for adapting to emerging modeling approaches, supporting collaboration across structural biology, computational biology, and related fields.

The impact of this work extends beyond model accuracy. With standardized, customizable datasets, CryoDataBot enables transparent benchmarking and cross-validation of AI models. Such transparency may help establish best practices in AI-driven structural biology, fostering a collaborative environment where models can be fairly compared on a consistent data foundation. Moreover, CryoDataBot’s adaptability ensures its continued relevance as cryoEM and AI technologies advance, positioning it as a vital tool in the field for years to come.

## Methods

### Retrieving Metadata for EMDB Entries

Our tool searches the EMDB for entries based on user-defined keywords, such as molecule type, organism, and resolution range. In the Electron Microscopy Data Bank (EMDB), each deposited cryoEM map is assigned a unique EMDB ID (e.g., “EMD-1234”). These cryoEM maps are often associated with corresponding atomic models, which have been fitted to the cryoEM maps. The atomic models are deposited separately in the Protein Data Bank (PDB) and are identified by their own PDB IDs (e.g., “6XYZ” or “7ABC”). CryoDataBot automatically retrieves a list of EMDB IDs and their associated fitted PDB IDs along with other metadata. If multiple PDB IDs are associated with an EMDB entry, only the first PDB ID is retained for downstream processing. Key metadata fields include: EMDB ID, fitted PDB IDs, the entry title, resolution (in Å), UniProtKB cross-references, AlphaFold cross-references, Q-score (a quantitative measure assessing atom resolvability in cryoEM maps), atom inclusion (the percentage of fitted atoms present in the density map), and the recommended contour level (the optimal threshold for visualizing electron density in 3D reconstructions). The ribosome dataset was retrieved on September 11, 2024, and the G protein dataset on August 14, 2025, to support reproducibility.

### Two-Tier Redundancy Filtering

Certain entries may represent the same biomacromolecule, though they are not exact duplicates. A biomacromolecule may have multiple EMDB entries due to variations in its conformational states or its binding to different ligands. To mitigate the overrepresentation of identical biomacromolecules or redundant structures within similar biomacromolecules, CryoDataBot employs a two-tier redundancy filtering process.

To assess the similarity between entry pairs, CryoDataBot leverages UniProtKB and AlphaFold cross-references, which correspond to functionally annotated protein domains and their respective structural predictions. Structural redundancy between pairs is quantified based on the proportion of shared cross-references, with a greater overlap signifying a higher degree of shared protein domains and, consequently, greater structural similarity.

The first-tier filtering, uniqueness filtering, removes entries with entirely overlapping cross-references, ensuring that only the highest-resolution entry is retained for each redundant group. Furthermore, uniqueness filtering flags and removes entries lacking both UniProtKB and AlphaFold cross-references. These flagged entries are stored separately for optional manual review, with the option to re-add them if deemed useful.

The second-tier filtering, similarity filtering, offers users the option to customize the desired level of dataset redundancy. Similarity filtering discards entries where the overlap ratio of cross-references exceeds a user-defined threshold, while retaining the entry with the highest resolution within each redundant group. This approach provides flexible control over dataset redundancy, allowing users to tailor the redundancy level to their specific needs, as the degree of redundancy directly influences the dataset size.

### Customized Map Resampling and Adaptive Map Normalization

The methodology in this study provides a more efficient and flexible approach to cryoEM map conditioning. While previous methods (e.g., DeepTracer^12^, Cryo2StructData^25^) rely on ChimeraX^11^ for map resampling, CryoDataBot leverages the *cupyx.scipy.ndimage.zoom* function from the CuPy Python library. This tool, integrated into the overall Python code, enables the resampling of cryoEM maps into a customizable uniform voxel size, with a default configuration of 1.0 Å × 1.0 Å × 1.0 Å. It eliminates the dependency on ChimeraX, thereby improving both the efficiency and flexibility of the resampling process.

In addition, our normalization strategy for cryoEM map densities addresses key limitations found in previous methods. Previous studies typically apply a fixed denoising threshold of zero: density values below zero are eliminated, positive values are retained, and the remaining values are normalized to the 0–1 range^12,14,25^. While this method removes low-intensity background values (noise) that do not correspond to molecular signal—defined here as high-density regions associated with structural features—it fails to account for variability in value ranges and noise-signal boundaries across different cryoEM maps. As a result, the fixed-threshold approach may lead to suboptimal denoising and normalization, particularly for maps with non-standard density distributions.

To address these limitations, we introduce an adaptive thresholding method tailored to the unique characteristics of each individual map. Rather than applying a fixed threshold of zero, our approach utilizes the recommended contour level—a metadata-defined reference value that typically exceeds zero and indicates the density level above which structural features are likely to be meaningful and visually interpretable. We compute a map-specific denoising threshold such that the recommended contour level aligns with the 85th percentile of retained density values. All values below this threshold are discarded, while those above are preserved and normalized to the 0–1 range. This effective removal of low-intensity noise, including values between zero and the new threshold.

By adapting the threshold to each map’s density distribution, our method enhances the precision of denoising and improves the flexibility of normalization. Compared to previous fixed-threshold approaches, it offers more consistent performance across diverse datasets.

### Map–model Fitness (MMF) Validation

While Q-score provides a useful residue-level assessment of model quality, it is insufficient for detecting global inconsistencies between cryoEM maps and their corresponding atomic models. In particular, it often fails to filter out entries showing localized agreement but poor overall alignment—such as cases where large regions of the cryoEM map remain unmodeled, or substantial portions of the model are unsupported by density (see the MMF validation example in Fig. 1c and Fig. S1). To address these limitations, we implemented an additional validation step, termed MMF validation, following map and model download, to more thoroughly evaluate global structural consistency.

To quantify global map–model alignment, we compute a Volume Overlap Fraction (VOF) score as the primary fitness metric. After map normalization, both the cryoEM map and the atomic model are projected into 2D along six directions—the three principal axes (X, Y, Z) and three diagonal-like orientations. Each projection is obtained by summing voxel values along the axis orthogonal to the projection plane (e.g., summing along the Z-axis for the XY projection). The resulting 2D images are then binarized by setting all values ≥ 1 to 1 and values < 1 to 0. For each projection pair, we calculate the intersection-over-union (IoU), or Jaccard index^33^, defined as the ratio of overlapping pixels to the total number of unique pixels in both projections. The final VOF score is computed as the average IoU across all six projections, excluding the highest-scoring projection to mitigate directional bias. Entries with VOF scores below a user-defined threshold (ranging from 0 to 1, with 1 indicating perfect alignment) are discarded (Fig. S1).

As a secondary metric, we also compute a Dice-like coefficient for each projection, defined as the pixel overlap divided by the total number of pixels across both projections (i.e., a Dice coefficient without the factor of two). The average Dice-like score is reported alongside the VOF score to provide complementary insight into map–model consistency.

### Generating Structural Labels

Structural labels are generated from atomic models. A 3D voxel array is initialized to match the spatial dimensions of the corresponding resampled cryoEM map and is then superimposed with the atomic model. Users specify label values (e.g., 1, 2, 3, 4) for atomic groups of interest, such as ligands, secondary structures, amino acids, or individual atoms.

The atomic model is traversed chain by chain to extract the coordinates of the specified atomic groups. These coordinates are converted to voxel indices within the voxel array. For each target voxel, the algorithm computes the Euclidean distance to nearby atomic coordinates and assigns the voxel the label of any atom that falls within a user-defined labeling radius. If a voxel lies within the radius of multiple atoms, the label of the closest atom is assigned. ach voxel is assigned a label corresponding to a single atom.

Users can adjust the labeling radius (default 1.5 Å) to control how far atomic influence extends. There are two main factors that should be considered. First, the type of feature they want to label. For example, labeling a single atom may require only a radius slightly larger than the atom itself, while larger features such as secondary structures can tolerate a bigger radius. Second, the tradeoff between maximizing positive labels and avoiding overlapping. In general, it’s best to select the largest radius that does not cause neighboring features to overlap. If some overlap is acceptable (as with secondary structures), the radius can be increased accordingly. For secondary structures and in general larger features, the optimal value may require some trial and error, but since labeling is a quick process, it can be repeated with ease.

After labeling is finished, the labeled volume is divided into smaller cubic sub-volumes (grids), with each grid saved as an individual .npy file. The grid size (default 64³ voxels) should be chosen based on the user’s training requirements. Most importantly, it should match the input size expected by their model. Beyond that, the only constraint is practicality: avoid excessively large values (e.g., 300^3^ or more), which may exceed the dimensions of some maps.

### Training the U-Net model

As shown in Figure 3a, we trained two independent 19-layer 3D U-Net models to predict secondary structures from cryoEM maps. Each model takes a 64³ Å³ density sub-volume as input and outputs N=5 probability volumes of the same size, representing voxel-wise probabilities for 5 classes: *Nothing*, *Helix*, *Sheet*, *Coil*, and *RNA*. Both models were trained under identical settings using a batch size of 16, the Adam optimizer^34^ with a learning rate of 1 × 10^−4^, and a weighted cross entropy loss to address class imbalance, with class weights set to [1, 57, 128, 52, 22]. Training was performed on an NVIDIA GeForce RTX 4090 GPU with 24 GB of display memory. We applied early stopping with a patience of 30 epochs, leading to training durations of 9.8 days (162 epochs) and 2.7 days (130 epochs) on the control and experimental datasets, respectively. The best-performing checkpoints were selected at epoch 132 for the control model and epoch 94 for the experimental model. Corresponding training and validation loss curves are shown in Figure 3b.

For the U-Net training on the G protein dataset, we used the same settings but with a batch size of 8. The U-Net was trained to identify C-alpha positions, producing N=2 probability volumes of 64³ Å³, representing voxel-wise probabilities for *Nothing* and *C-alpha*. The class weights for the weighted cross-entropy loss were assigned based on the number of samples in each class. This resulted in the best-performing checkpoints being selected at epoch 111, 68, 63, and 43 for the raw, Q0.3, experimental (Q0.4) and Q0.5 model, respectively. Training and validation loss curves are shown in Figure 7a.

### Evaluating AI model prediction performance

The outputs of both the U-Net model and CryoREAD Stage 1 are voxel-based, with each voxel assigned a probability distribution over all structural labels. For example, in the U-Net model predictions, each voxel contains a probability vector ***p*** = (*p_0_*, *p_1_*, *p_2_*, *p_3_*, *p_4_*), representing the probabilities of the voxel belonging to one of five classes: 0, 1, 2, 3, 4, corresponding to *Nothing*, *Helix*, *Sheet*, *Coil*, and *RNA*, respectively. For evaluation, a voxel is assigned to label *k* (*k* ∈ {1, 2, 3, 4}) if *pₖ*exceeds 0.8 (or 0.4 for CryoREAD Stage 1, as recommended by its first author); if no probability exceed the threshold, the voxel is assigned to label 0 (*Nothing*). Ground truth labels were generated from deposited atomic models using the same procedure described in the **Generating Structural Labels** section.

To evaluate prediction performance for each label, we computed voxel-wise accuracy, precision, recall, and F1 score. For a given label *k*, TP, FP, FN, and TN denote the number of voxels that are correctly predicted as *k*, incorrectly predicted as *k*, missed ground truth voxels of *k*, and correctly predicted as not *k*, respectively. The evaluation metrics are calculated as follows:

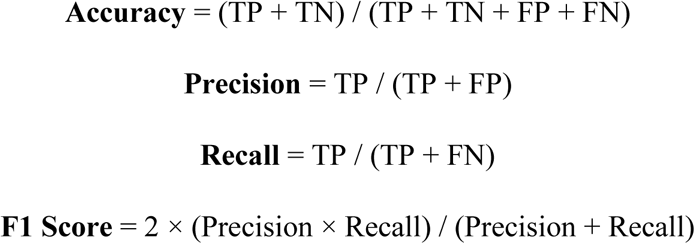

Accuracy measures the proportion of all predicted labels, across all classes, that match the ground truth. Precision quantifies the fraction of correct predictions among all predicted labels, reflecting the reliability of the model’s outputs. Recall captures the proportion of ground truth labels that were correctly identified, indicating the model’s sensitivity to relevant features. The F1 score, defined as the harmonic mean of precision and recall, provides a balanced evaluation of both correctness and coverage—particularly useful in cases of class imbalance or when precision and recall diverge. These metrics were computed separately for each structural label and in aggregate, both during and after training, to enable detailed performance assessment.

### Graphical User Interface (GUI) for the Tool

To improve usability, we developed a user-friendly GUI using PyQt5, a Python binding for the cross-platform Qt framework. The interface (Fig. 1e) features an intuitive design that streamlines data retrieval, quality control, and label selection, thereby facilitating efficient user interaction with the tool.

For data retrieval, users can formulate queries using EMDB’s native search syntax^28^. The application then fetches and downloads a comma-separated values (CSV) file containing the corresponding entries and associated metadata. For quality control, users may proceed with the retrieved CSV file or upload a custom CSV file, followed by the specification of filtering parameters to curate the dataset. For label selection, the interface allows users to define atomic groups by configuring options across three hierarchical categories: the **secondary structure** category includes –*helix*, *sheet*, *coil*, *RNA*, *and DNA*; the **residue type** category supports the 20 standard amino acids and the 4 canonical nucleobases; and the **atom type** category allows input of specific atom names in PDB files (e.g., *CA*, *Mg*). For instance, to label all Cα atoms within helices, the user would select “helix” under secondary structure, omit residue type selection, and input “CA” as the atom type. To label other structures (e.g., a ligand), the user would select “none” under secondary structure, leave the residue type field blank, and input all relevant atom names under atom type (e.g., the ligand or its constituent atoms).

## Data Availability

The entries of the cryoEM maps and their corresponding atomic models used in this study are listed in the Supplementary Tables. The cryoEM maps can be downloaded from the EMDB via the European Molecular Biology Laboratory – European Bioinformatics Institute (EMBL-EBI) FTP server (https://ftp.ebi.ac.uk/pub/databases/emdb/structures). The corresponding atomic models are available from the Research Collaboratory for Structural Bioinformatics (RCSB, https://www.rcsb.org/). All data that support this study are available from the corresponding authors upon request.

## Code Availability

The source code of CryoDataBot is available at https://github.com/t00shadow/CryoDataBot under the MIT license. CryoDataBot has been registered in WorkflowHub^35^. Its Research Resource Identifier (RRID) is *SCR_027186*, and its biotoolsID is biotools:cryodatabot (https://bio.tools/cryoDataBot).

## Contributions

Z. Hong Zhou conceived and oversaw the project, participated in project design and result illustration, and wrote the paper. Qibo Xu co-conceived the project and participated in all aspects of its development. Leon Wu designed and built the GUI, assisted with debugging, optimizing, and organizing the backend code, prepared Figures 1e and 3b, and drafted the Methods section. Michael Rebelo developed the redundancy evaluation method, contributed to backend debugging and optimization, and assisted with manuscript writing. Shi Feng implemented the Linux command-line interface, optimized the customized dataset construction module, contributed to the metadata collection and structural data conditioning modules as well as the redundancy evaluation method, prepared Figures 2b and 2c, and drafted the Methods section. Xinye Yu developed the Q-score fetching function, helped test and optimize the software, assisted in creating Figures 4 and 6, and drafted the Discussion section. Farhanaz Farheen retrained CryoREAD, evaluated the results, helped prepare Figure 5, and drafted the first paragraph of the CryoREAD retraining section. Daisuke Kihara supervised the CryoREAD retraining and evaluated the results. All authors edited and approved the manuscript.

## Acknowledgements

This project is supported by a grant from the US National Institutes of Health (R01GM071940 to Z.H.Z.). D.K. acknowledges supports from the NIH (R01GM133840) and the National Science Foundation (IIS2211598).

We thank Xiao Wang for technical help regarding training and evaluation of the retrained CryoREAD model.

---

**Fig. S1.**
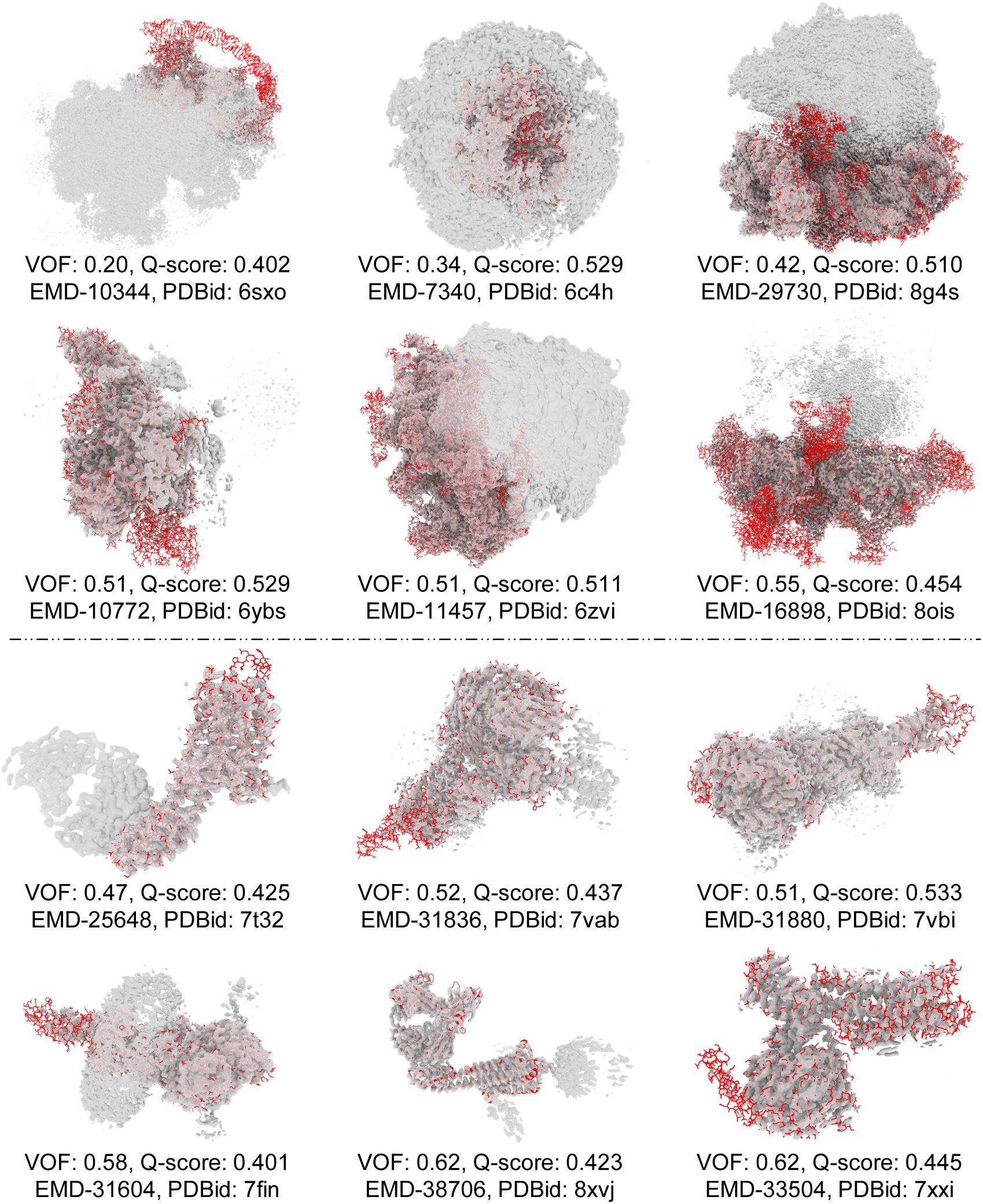
Examples discarded during MMF validation (low VOF score despite good Q-score). The top six examples are selected from 71 entries excluded during the construction of the ribosome experimental dataset, while the bottom six are drawn from 6 entries excluded during the construction of the G protein experimental dataset. The VOF score effectively captures global inconsistencies that may not be detected by Q-score alone.

**Fig. S2.**
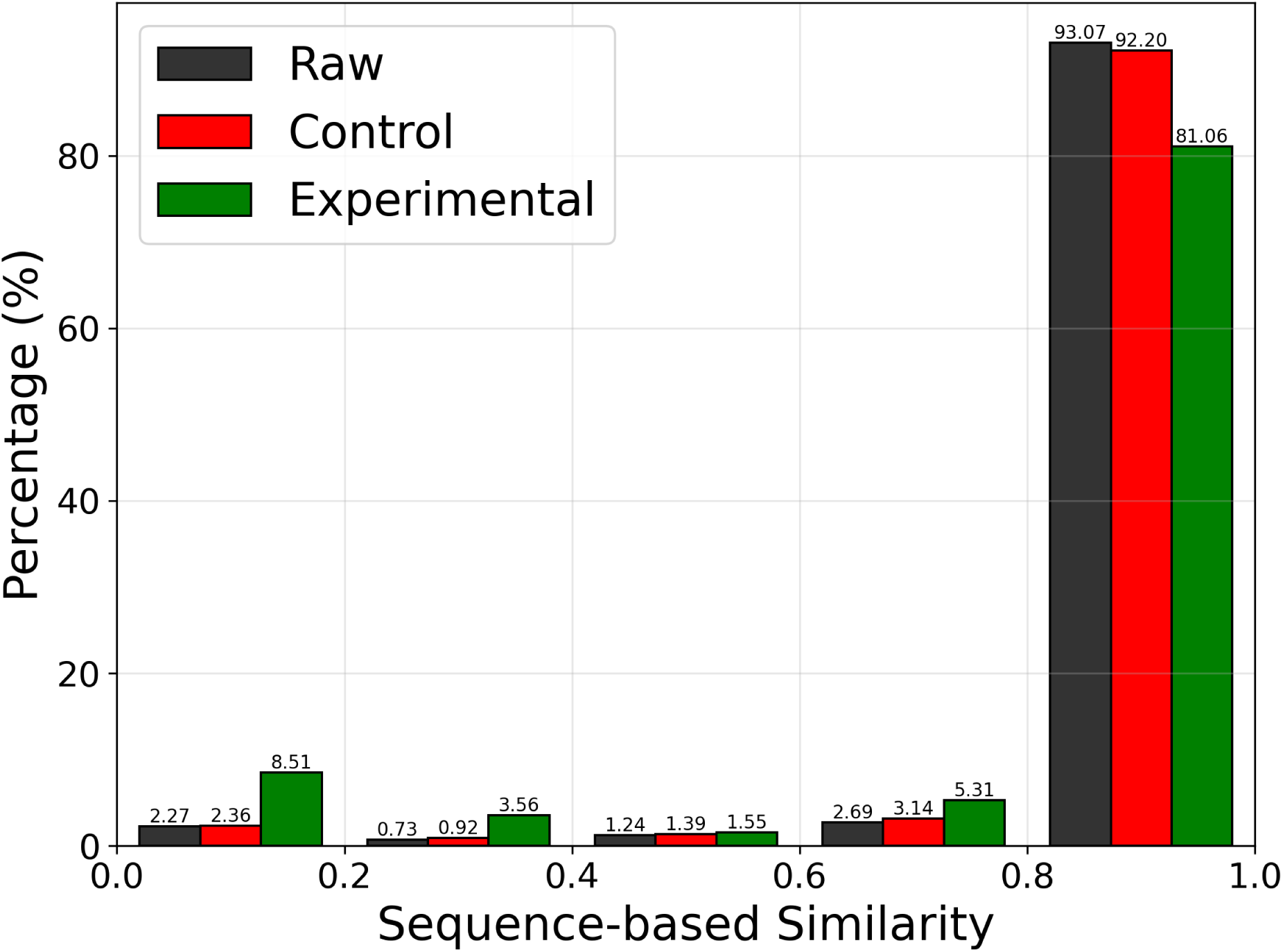
Comparative analysis of sequence similarity across datasets. For each entry **X** (containing *x* chains), each chain was compared via BLASTp against every chain from all other entries. If another entry **Y** (containing *y* chains) has *n* chains identified as homologous (e-value < 0.001 from the BLASTp results) to any chains in **X**, the sequence-based similarity **Y** to **X** is defined as *n/y*. This asymmetric scoring yields *N*(N–1)* pairwise similarity scores for a dataset with *N* entries. A larger proportion of high scores indicates greater sequence-level redundancy. As shown, the Raw and Control datasets exhibit comparable redundancy, whereas the Experimental dataset shows a marked reduction—consistent with the structural similarity trends in Fig. 2b.

**Fig. S3.**
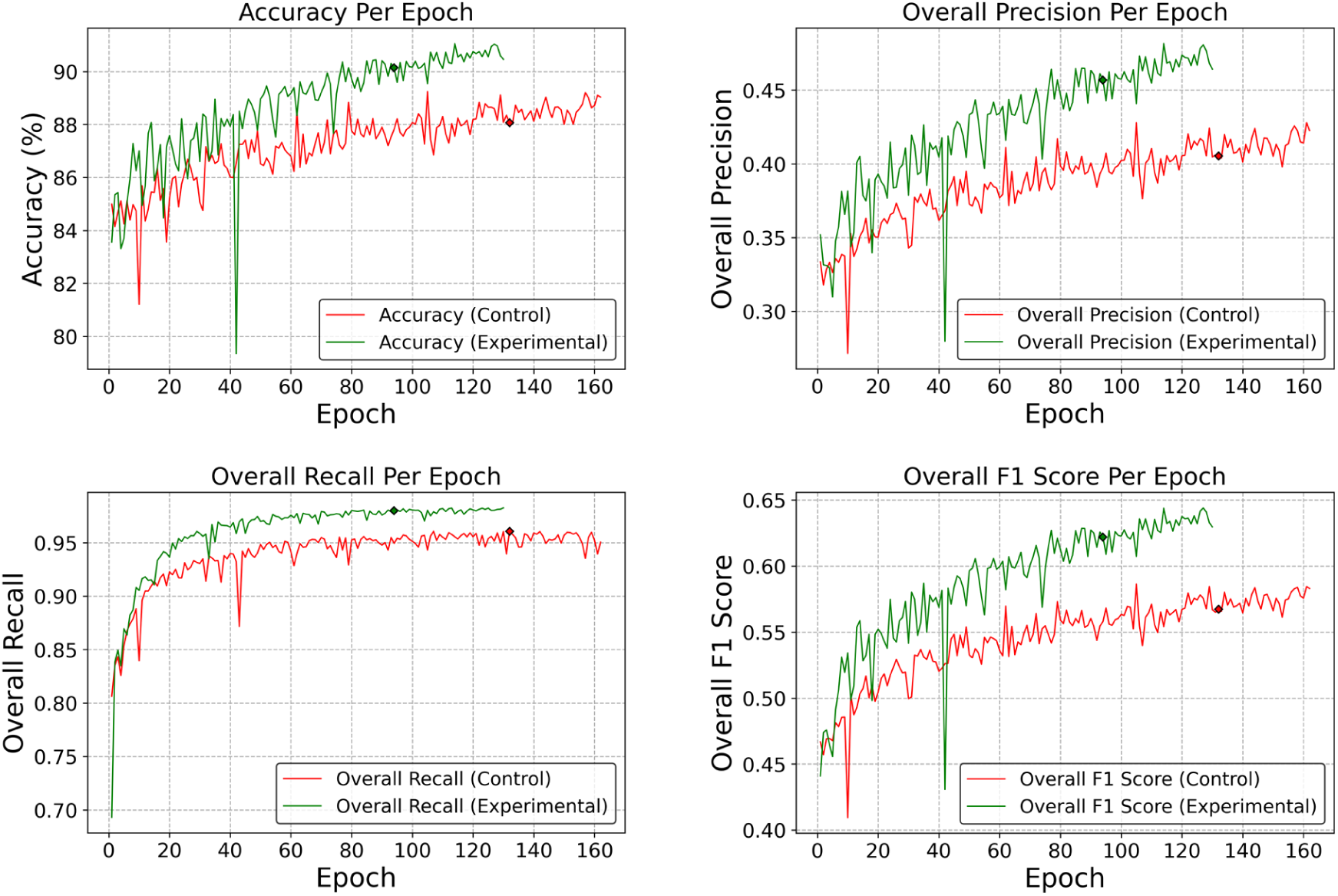
Training dynamics of the U-Net models on control and experimental datasets. Evolution of overall accuracy, precision, recall, and F1 score on the validation set during training of the control model (red) and the experimental model (green). Epochs corresponding to peak performance are marked by diamonds at epoch 132 (control) and epoch 94 (experimental). The experimental model achieved higher validation metrics in fewer epochs, indicating improved training efficiency.

**Fig. S4.**
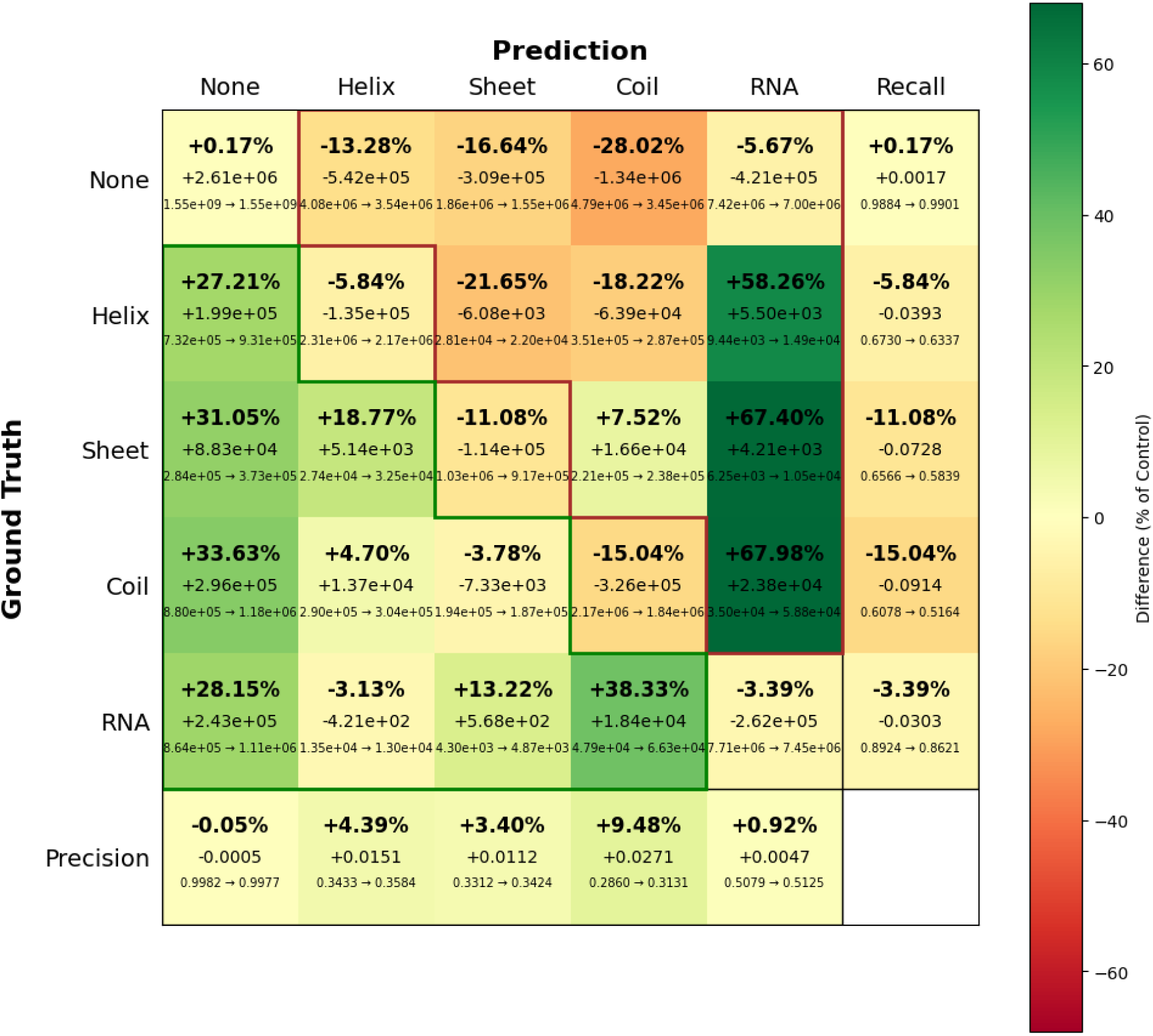
Comparison of confusion matrices between the control and experimental models for secondary structure prediction, including recall and precision metrics. Each cell displays three lines: (bottom) control model count → experimental model count, (middle) the absolute difference (experimental minus control), and (top) the ratio of difference to control. Cells are color-coded based on this ratio. Brown outlines indicate false positives (FP), relevant to precision calculation, while green outlines indicate false negatives (FN), relevant to recall calculation.

**Fig. S5.**
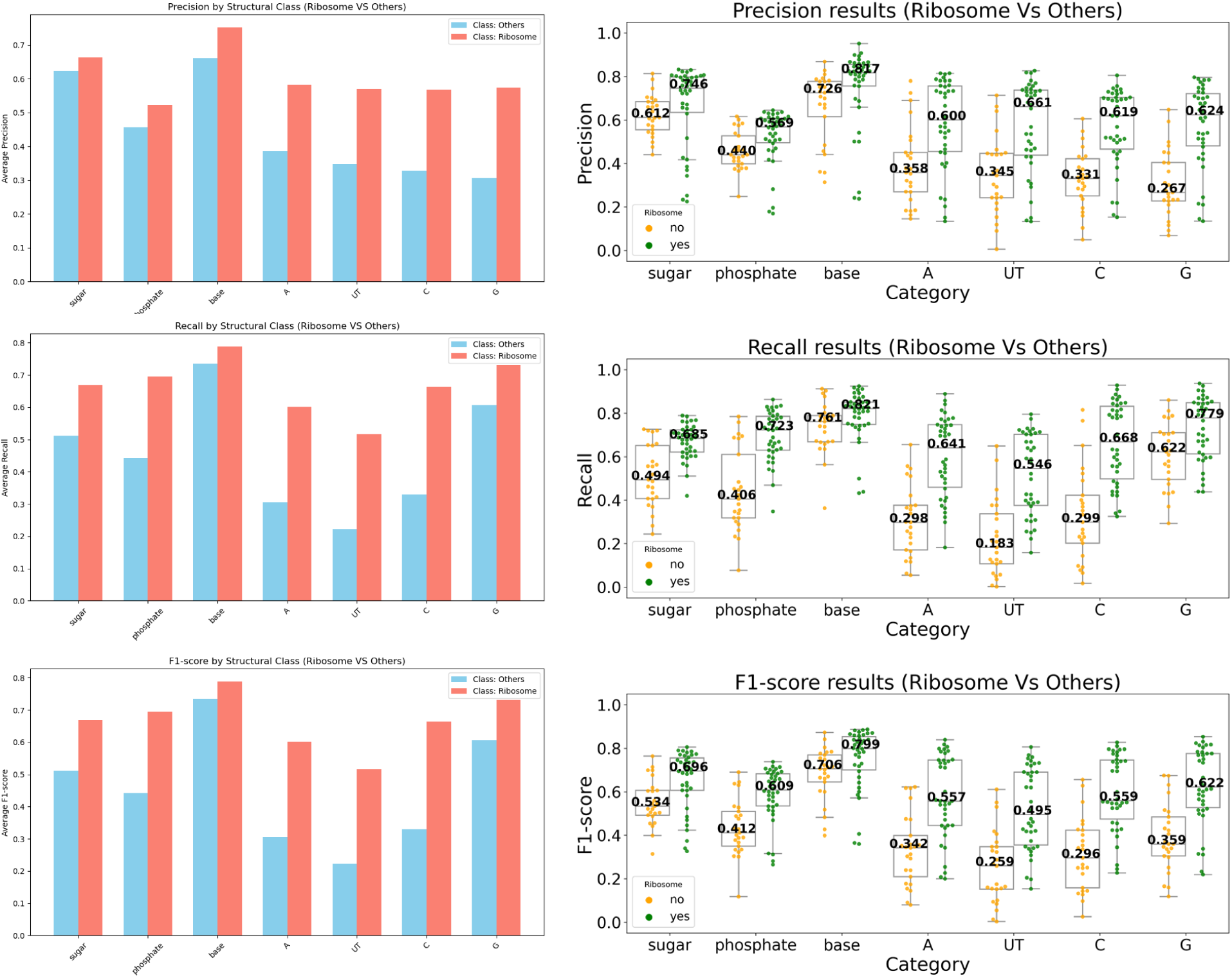
Comparison of Precision, Recall and F1-scores for retrained CryoREAD with CryoDataBot training set showing the difference in performance on ribosomes and other classes in the test set. The bar charts show average precision, recall, and F1 scores for ribosomes (38 cases) and other classes (25 cases). The box plots show the individual data points representing EMDB maps along with the medians annotated. Since the retrained CryoREAD with CryoDataBot dataset was trained with exclusively ribosome-based data, the performance on the test set is affected by it. For all 7 structural classes (Sugar, Phosphate, Base, A, U/T, C, G) the metric values for ribosome test cases are higher than those on other classes.

**Fig. S6.**
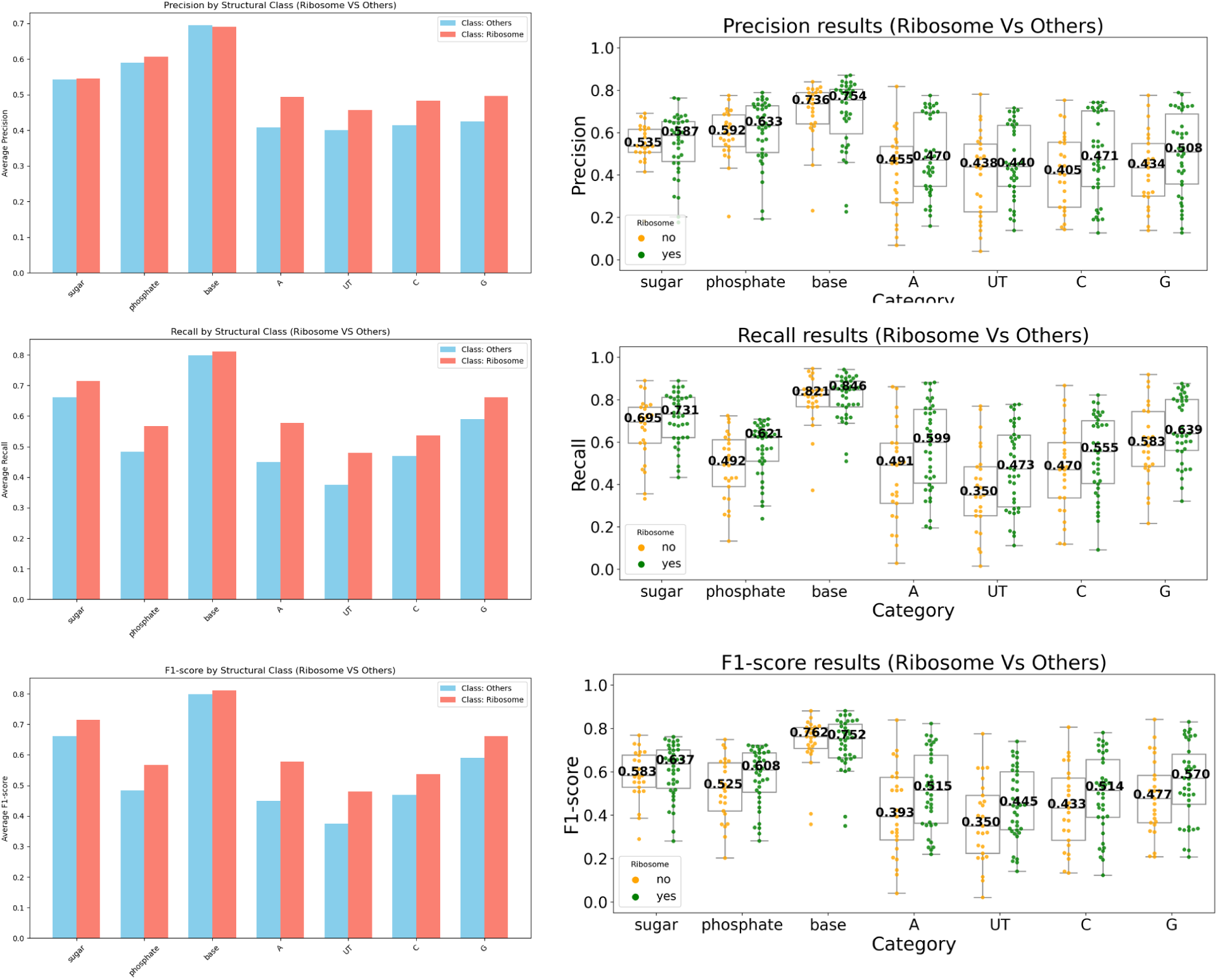
Comparison of Precision, Recall and F1-scores for original CryoREAD showing the difference in performance on ribosomes and other classes in the test set. The bar charts show average precision, recall, and F1 scores for ribosomes (38 cases) and other classes (25 cases). The box plots show the individual data points representing EMDB maps along with the medians annotated. The original CryoREAD was trained with different classes of RNA and DNA. Contrary to Fig. S5, the difference in performance for ribosome test sets and other classes of RNA/DNA is less. The average base precision is higher for other RNA/DNA classes than for ribosomes. The box plot for F1-scores also shows that the median F1-score for other classes is higher than those with ribosome test cases.

**Fig. S7.**
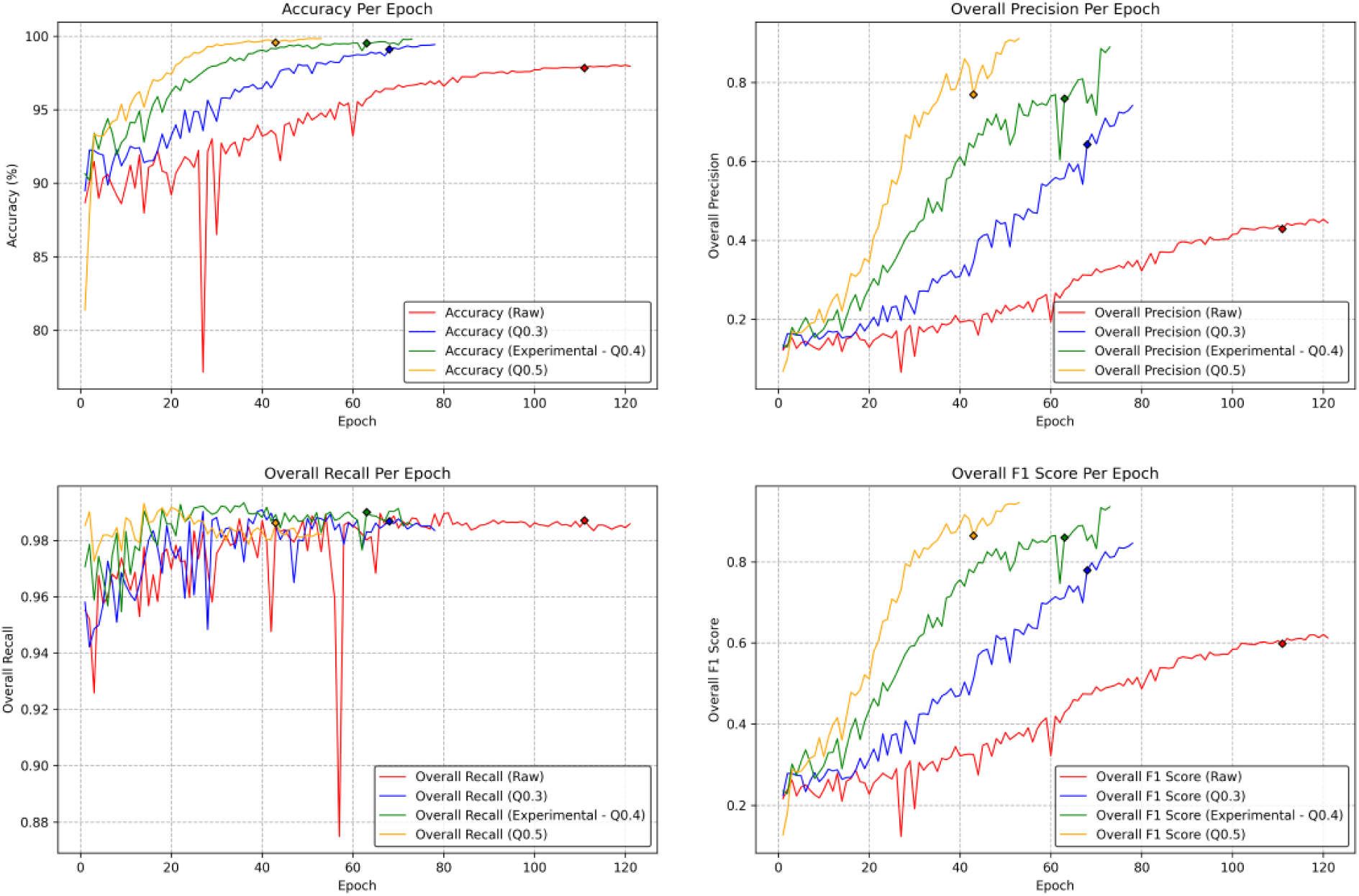
Training dynamics of U-Net models training on G protein datasets. Evolution of overall accuracy, precision, recall, and F1 score on the validation set during training of the raw (red), Q0.3 (blue), experimental (green) and Q0.5 (orange) models. Epochs corresponding to the best performance are marked by diamonds at epoch 111 (raw), 68 (Q0.3), 63 (experimental - Q0.4), and 43 (Q0.5). Compared to the raw model, the experimental model reached higher validation metrics in fewer epochs, indicating improved training efficiency.

